# Biased localization of actin binding proteins by actin filament conformation

**DOI:** 10.1101/2020.02.21.959791

**Authors:** Andrew R Harris, Pamela Jreij, Brian Belardi, Andreas Bausch, Daniel A Fletcher

## Abstract

The assembly of actin filaments into distinct cytoskeletal structures plays a critical role in cell physiology, but how proteins localize differentially to these structures within a shared cytoplasm remains unclear. Here, we show that the actin-binding domains of accessory proteins can be sensitive to filament conformational changes. Using a combination of live cell imaging and *in vitro* single molecule binding measurements, we show that tandem calponin homology domains (CH1-CH2) can be mutated to preferentially bind actin networks at the front or rear of motile cells, and we demonstrate that the affinity of CH1-CH2 domain mutants varies as actin filament conformation is altered by perturbations that include stabilizing drugs, physical constraints, and other binding proteins. These findings suggest that conformational heterogeneity of actin filaments in cells could help to direct accessory binding proteins to different actin cytoskeletal structures through a biophysical feedback loop.

## INTRODUCTION

Multiple actin cytoskeletal structures co-exist within the cytoplasm, yet they are spatially organized, architecturally distinct, and perform specific functions^1,2^. In addition to branched actin networks in the lamellipodium, and stress fibers in the cell body, advances in both optical and electron microscopy continue to reveal more details about the organization and assembly a broader range of actin structures, including filopodia^3^, asters and stars^4^, podosomes^5^ and patches^6^. In each of these structures, the interaction of actin filaments with a vast set of accessory proteins promotes the formation of distinct cytoskeletal architectures.

Interestingly, common probes for f-actin, including GFP-tagged actin, small actin-binding peptides (lifeact^7^, f-tractin^8^, affimers^9^) and phallotoxins^10^, are known to not distribute evenly on different actin cytoskeletal structures^11–13^. Similar observations have been made for fluorescent fusions to minimal actin binding domains from different proteins^14–17^. Mechanistically, these results have been attributed to the complex and competitive interactions between side binding proteins^18,19^, the effect of actin nucleators^20,21^, and the kinetic properties of the reporting probe^10,22^. However, other properties of actin filaments, including its conformational state, could differ among cytoskeletal structures due to biophysical constraints and be detected by actin-binding proteins to bias their localization.

Several studies have indicated that the conformational state of an actin filament is polymorphic^23–25^ and that actin filaments exist in a range of different ‘flavors’, including different their nucleotide state^26^, oxidative state^27^ and twisted states^23,28,29^. Actin binding proteins have also been shown to modulate filament structural conformations either as part of their regulatory activity or as a means for allosteric cooperative binding to actin^30–32^. In addition to effects of protein binding, mechanical perturbations to actin filaments such as torques, tension^33,34^ and bending have been suggested to influence protein interactions with filaments, including the binding of the Arp2/3 complex^21^ and severing activity of the protein cofilin^20,35,36^. Together, these observations suggest that different conformations of f-actin, induced either mechanically or biochemically, could impact the affinity of actin binding proteins for f-actin.

We sought to investigate whether filament conformational changes could be sensed by a common class of actin binding domain, tandem calponin homology domains (CH1-CH2), and whether affinity changes for different conformations could be responsible for biasing the localization of CH1-CH2-containing proteins to different actin structures in cells. Using a combination of live cell imaging and *in vitro* single molecule binding affinity measurements, we find that mutants of the actin-binding domain of utrophin (CH1-CH2) localize to different actin structures and exhibit differing binding affinities for actin filaments whose conformational state has been altered biophysically and biochemically. We also show that this mechanism extends to native actin-binding domains, suggesting that sensitivity to actin filament conformational states could be playing an important role in the organization and regulation of actin binding proteins in actin filament structures.

## RESULTS

### Utrophin actin binding domain mutants localize to different actin structures

CH1-CH2 domains are found in many actin crosslinking and regulatory proteins^37^ including α-actinins in stress fibers^38^ and filamins in the actin cortex^39^. The actin binding domain of utrophin (CH1-CH2) is used as a generic marker for f-actin^40^, which raised the question of whether it uniformly labels all filaments in cells or might be sensitive to filament conformational heterogeneity. Recent cryo-Electron Microscopy studies have mapped the interacting residues between the actin binding domain from filamin A and f-actin^41^, and between the actin binding domain of utrophin and f-actin to 3.6Å resolution^42^. Three major regions on CH1 interact with f-actin, ABS-N, ABS2 and ABS2’, which make contacts with two longitudinally adjacent subunits in an actin protofilament^41,42^, suggesting that these domains might be sensitive to actin filament conformation. We chose to examine residues predicted to lie within actin binding surface 2 (ABS2) on utrophin CH1^41^, at the CH1-CH2 domain interface^37,43^ and within the ABS-N (also referred to as the n-terminal flanking region)^37,44,45^, which we had found altered actin-binding affinity in a previous study^37^.

We first compared the localization of binding interface mutations to that of the actin binding domain of utrophin (utrnWT) in both HeLa cells and PLB neutrophils, which have clearly distinct actin structures (Fig 1, Fig S1). Interestingly, we observed several combinations of mutations that caused significant changes in localization relative to utrnWT (see *Materials and Methods, Engineering utrn ABD affinity and specificity*). The mutations Q33A T36A K121A caused an increased enrichment to lamellipodial actin (Fig 1A, Movie S1, Movie S2), while Q33A T36A G125A L132A was comparatively enriched at the rear of the cell (Fig 1B). We previously showed that truncating the n-terminal region prior to CH1, Δ-nterm Q33A T36A, changes binding to focal adhesions in HeLa cells^37^ (Movie S3), and this mutant was more evenly distributed at the rear and front of migrating neutrophils compared to utrnWT (Fig 1C, Movie S4). Subsequently, we refer to the minimal actin binding domain of utrophin as utrnWT, Q33A T36A K121A mutant as utrnLAM, the Δ-nterm Q33A T36A mutant as utrnΔN, and the Q33A T36A G125A L132A mutant as utrnSF.

**Figure 1:**
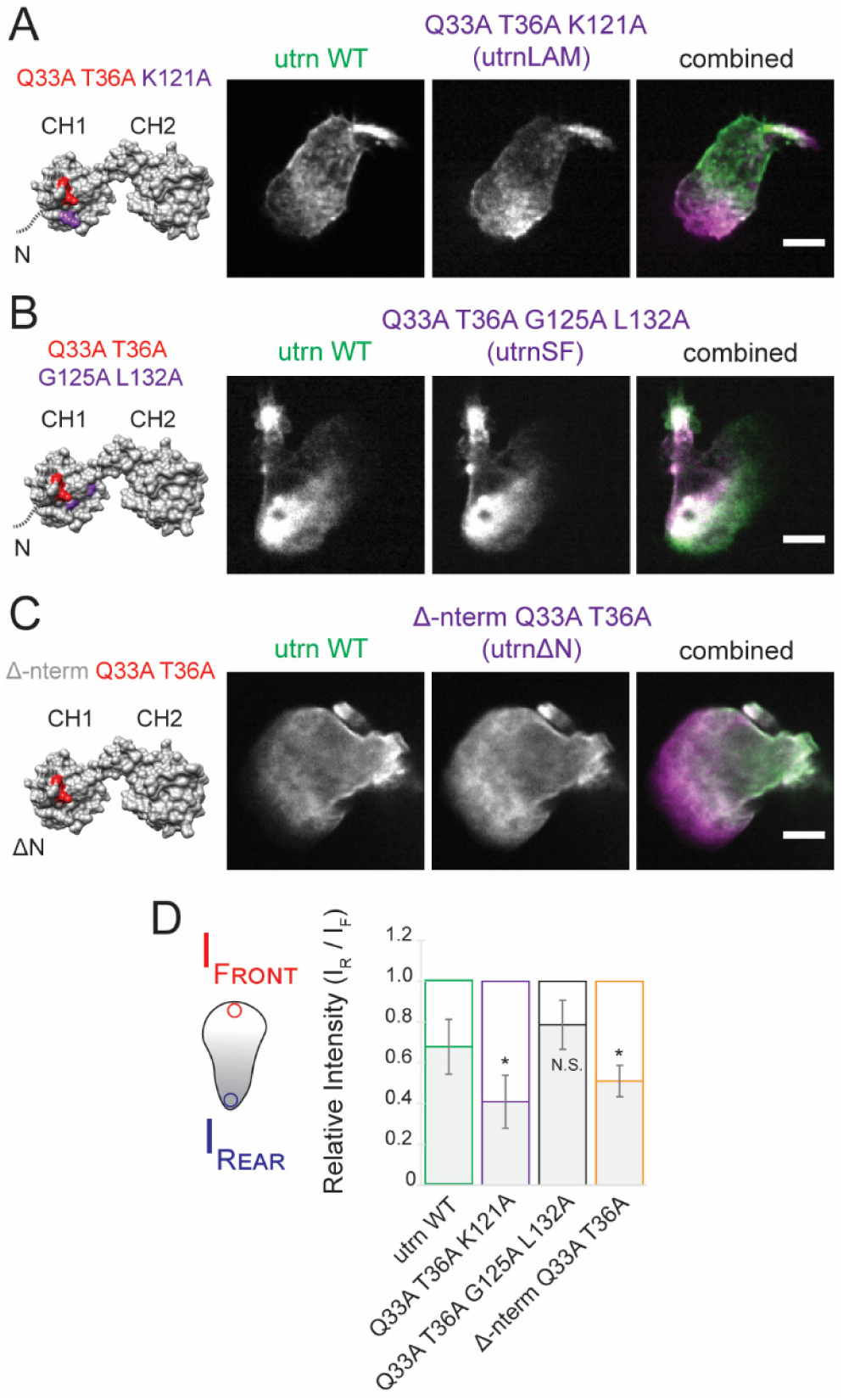
utrn CH1-CH2 mutants display differential front-back localization in neutrophils. Structure of the actin binding domain of utrophin (1QAG ^73^). The N-terminal flanking region which is not resolved in the unbound crystal structure (1QAG) becomes structured upon actin binding^38,39,42^ and is indicated by the grey line in the image. Images show the mutant in magenta compared to utrnWT in green. (A) The mutant utrn Q33A T36A K121A is localized more strongly to the leading edge than utrnWT. (B) The mutant utrn Q33A T36A G125A L132A is localized more strongly to the rear of the cell than utrnWT. (C) The mutant utrn Δ-nterm Q33A T36A is localized more strongly to the leading edge than utrnWT, similar to utrn Q33A T36A K121A. (D) Comparisons of the relative utrn construct intensity at the front and back of migrating neutrophils, calculated by averaging the intensity in 1µm regions at the front and back of the cell (left).

### Single molecule dwell times report complex behavior of utrophin mutants

We wondered whether the differences in localization of these domains could arise due to a bias in binding affinity for actin filaments in each specific network – which we refer to as specificity. We sought to characterise the binding properties of each mutant in more detail *in vitro*. Previously, single molecule kinetic measurements have been used to investigate the binding properties of actin severing proteins^46^, formins^47^, cofilin^48^ and the actin binding domain of α-catenin^49^. For α-catenin, the binding dwell time of single molecules (inverse of the off-rate) was shown to follow a two-timescale binding behaviour, in which the binding dwell times increase as a function of concentration of the domain added. This cooperative change in dwell time was hypothesized to be due to structural changes in f-actin that are induced by α-catenin’s actin binding domain binding to f-actin^49^. Dynamic chances in actin filament conformation in response to biochemical perturbations have also been measured using single molecule FRET measurements on dual-labelled actin monomers^24^. Therefore, to obtain a detailed understanding of the actin-binding kinetics of our different mutants and potential effects of actin structural conformation on binding, we used a TIRF-based single molecule binding assay to measure binding dwell times and binding rates (Fig 2A, *Materials and Methods, In vitro single molecule binding kinetics assay*).

**Figure 2:**
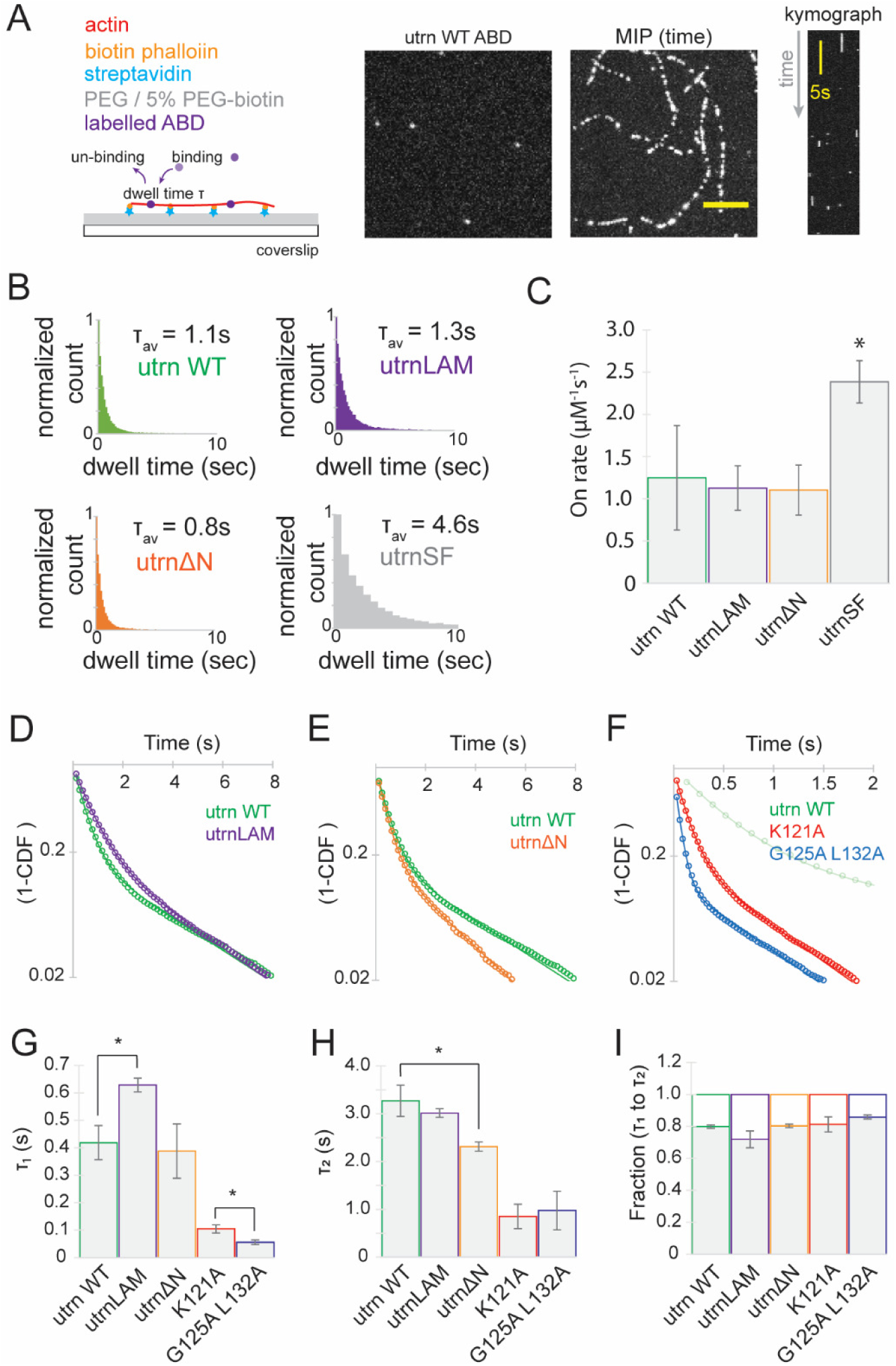
Single molecule kinetic measurements of utrn CH1-CH2 mutants in vitro. (A) Single molecule binding assay to measure the kinetic properties. Images in the example shown are for utrnWT. Maximum intensity projection through time displays the filament backbones and a kymograph the kinetics of binding. (B) Average binding dwell times for the different CH1-CH2 mutants. (C) Binding on rates for the different utrophin mutants. (D) Cumulative distribution function for utrnWT and utrnLAM. (E) Cumulative distribution function for utrnWT and utrnΔN. (F) Cumulative distribution function for K121A and G125A L132A. (G) Comparisons of the first timescale τ_1_, (H) second timescale τ_2_, and (I) relative amplitudes (a_1_, 1-a_1_) from a double exponential fit to the cumulative distribution functions for the different constructs.

We compared the distribution and lengths of binding dwell times for the different mutants identified in our first screen. The average dwell times were similar between utrnWT (τ_av utrnWT_ ∼ 1.1sec), utrnLAM (τ_av utrnLAM_ ∼ 1.3sec) and utrnΔN (τ_av utrnΔN_ ∼ 0.8sec) (Fig 2B). In contrast, utrnSF had a longer average dwell time (τ_av utrnSF_ ∼ 4.6sec), indicating that this mutant turned over more slowly. We measured the binding on-rate using our single molecule assay and found that utrnWT (k_on_ = 1.25 ± 0.36 µM^-1^s^-1^), utrnLAM (k_on_ = 1.13 ± 0.15 µM^-1^s^-1^), and utrnΔN (k_on_ = 1.10 ± 0.17 µM^-1^s^-1^) were similar and utrnSF (k_on_ = 2.38 ± 0.18 µM^-1^s^-1^) had a higher on rate (Fig 2C). Intrigued by the differences in localization and similar kinetics of utrnWT, utrnLAM and utrnΔN, we focused our attention on these mutants in particular.

For all of the mutants tested, the distribution of binding dwell-times was well characterised by a double exponential fit (R^2^=0.99 double exponential, R^2^= 0.94 single exponential, Pearson’s correlation coefficient), suggesting a two-timescale binding model best described the behaviour of these constructs (Fig 2D-F, Fig S3, *materials and methods* - *In vitro single molecule binding kinetics assay*). By comparison, the common actin binding probe Lifeact was well characterised by a single exponential (R^2^=0.99, single exponential, Fig S3 D), indicating that the mechanisms of binding for CH1-CH2 domains is more complex than that of the short peptide.

The cumulative distribution function (CDF) of binding dwell-times of the different mutants revealed further differences (Fig 2 D-F). The two-timescale behaviour of utrnWT (Fig 2D) and utrnΔN (Fig 2E) was more distinct than the flatter behaviour of utrnLAM, as characterised by a smaller difference in the two timescales (τ_2_/τ_1 utrnWT_ = 7.8, τ_2_/τ_1 utrnLAM_ = 4.8, τ_2_/τ_1 utrnΔN_ = 6.0) and a reduction in the relative amplitudes of the two timescales (a_1 utrnWT_ = 0.8, a_1 utrnLAM_ = 0.7, a_1 utrnΔN_ = 0.8) (Fig 2 G,H,I). Removing the Q33A T36A mutations from utrnSF and utrnLAM reduced the overall dwell-time of both (Fig 2F), suggesting that residues within ABS2 and the n-terminal flanking region were indeed important for direct interactions with actin and their localization.

### Filament stabilization by Jasplakinolide but not phalloidin alters utrophin ABD mutant dwell time

We next tested whether stabilization of actin filaments with the small molecules phalloidin and jasplakinolide which have been shown to have different effects on actin filament structural conformation^50^, altered the binding dwell-times of the different utrophin mutants (Fig 3A). We first introduced 1µM phalloidin into our single molecule binding assay after filaments were attached to the surface of the coverslip, and we found that it had no effect on the binding dwell-time of either utrnWT (τ_1 utrnWT_ = 0.42 ± 0.06 sec, τ_1 utrnWT+phall_ = 0.36 ± 0.03 sec p=0.14), utrnLAM (τ_1 utrnLAM_ = 0.63 ± 0.03 sec, τ_1 utrnLAM+phall_ = 0.52 ± 0.12 sec, p=0.50), or utrnΔN (τ_1 utrnΔN_ = 0.39 ± 0.01 sec, τ_1 utrnΔN+phall_ = 0.45 ± 0.04 sec, p=0.83) (Fig 3B-D, Fig S4 A). In contrast, introduction of 1µM of the actin stabilizing agent jasplakinolide affected both utrnWT and utrnLAM, making the dwell-time of single molecules shorter in both cases (τ_1 utrnWT+jasp_ = 0.27 ± 0.01 sec, p=0.05, τ_1 utrnLAM+jasp_ = 0.29 ± 0.02 sec, p<0.05, Fig 3B-D). Interestingly, the effect of jasplakinolide was stronger on utrnLAM (∼53% reduction in dwell time) than it was on utrnWT (∼36% reduction in dwell time), suggesting that each mutant had a different degree of specificity for jasplakinolide stabilized f-actin. Surprisingly, jasplakinolide treatment had little effect on the binding dwell time of utrnΔN (τ_1 utrnΔN+jasp_ = 0.43 ± 0.03 sec, p=0.09). This result suggests that this mutant was insensitive to actin filament conformational change induced by jasplakinolide.

**Figure 3:**
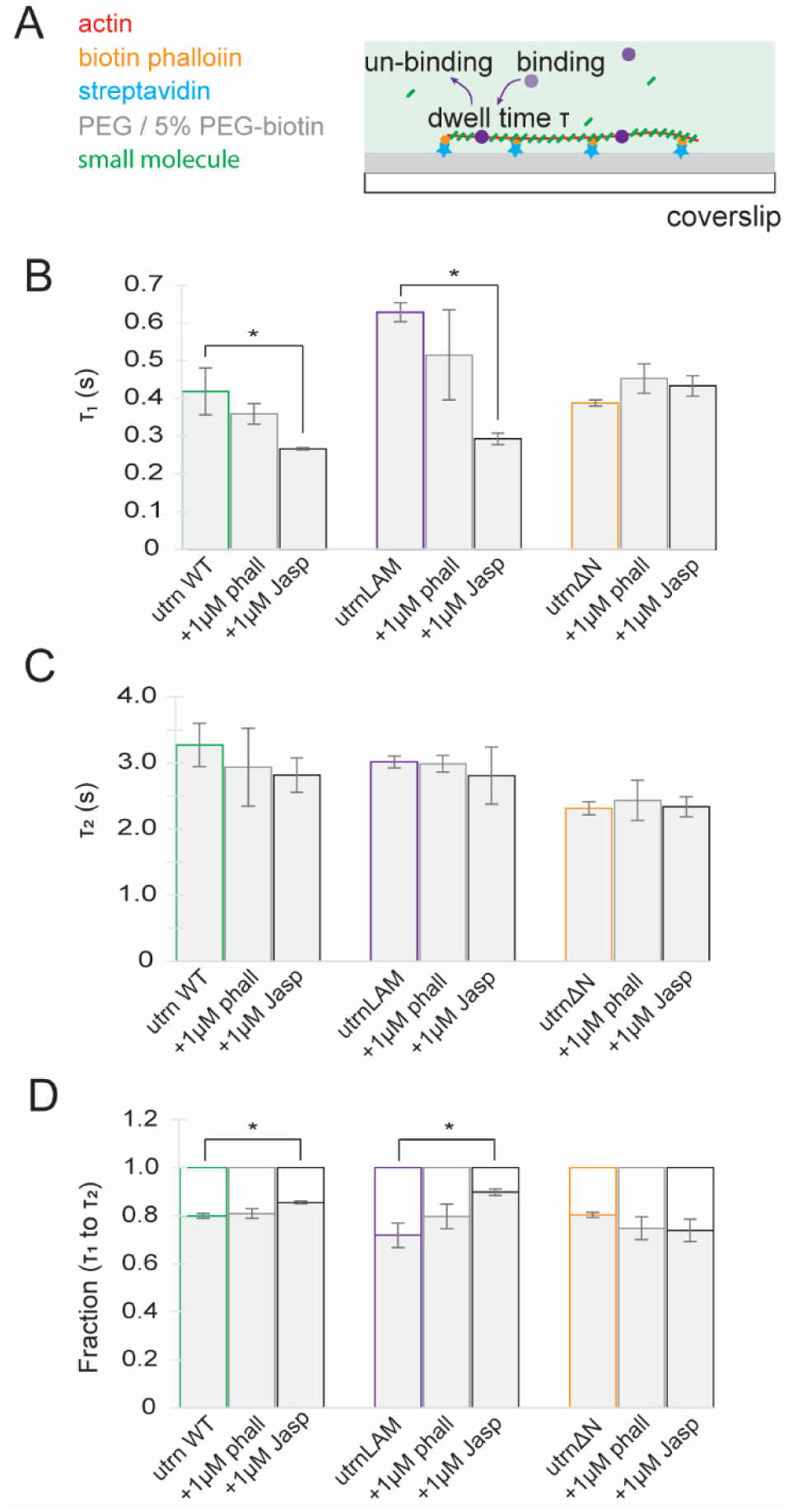
Filament stabilization by Jasplakinolide but not phalloidin alters utrophin ABD mutant dwell time. (A) Measurement of binding dwell times in the presence of actin stabilizing agents. (B) (B) Comparisons of the first timescale, (C) second timescale, and (D) relative amplitudes from a double exponential fit to the cumulative distribution functions for the different constructs and conditions.

### Filament binding by cofilin and drebrin alters dwell time of utrophin ABD mutants

In addition to the small molecules phalloidin and jasplankinolide, several actin binding proteins have been shown to impact filament structure. Cofilin is an actin severing protein that breaks actin filaments by forming discontinuities in filament mechanical properties^36,51^. Non-continuous mechanical properties are caused by local changes in filament twist induced by cofilin binding, which change the helical half pitch of f-actin from a mean of ∼36nm to ∼27nm^52^. Given our observations that utrophin ABD mutants were sensitive to actin filament conformation induced by jasplakinolide, we investigated how the different utrophin ABD mutants interacted with cofilin.

First, we measured the severing activity of cofilin in the presence of different utrophin mutants (Fig 4A). We found that introducing 200nM of each of the different mutants reduced the severing rate of 75nM cofilin^49^, likely due to direct competition for a similar binding site on f-actin (Fig 4B). Next, we sought to test whether conformational changes induced by cofilin binding impacted the dwell time of the different mutants. We used a dual-color binding assay with a low concentration of labelled cofilin (10nM), which is not sufficient to drive filament severing, and single molecule levels of utrophin mutant ABD. We then sorted the utrophin mutant ABD single molecule binding events based on their distance from cofilin clusters^48^, which we were able to localize with a precision of ±30nm (Fig 4C). Since structural changes in actin induced by cofilin are reported to propagate locally, distances ranging from 1-2 subunits^53,54^, we considered single molecule binding events within 30nm from a cofilin binding event to be ‘near’ and those beyond 30nm to be ‘far’. We measured generated the cumulative distribution functions for near and far cofilin molecules (Fig S4 D-F) and compared the τ values. Due to the low number of near events (∼1% of total events) we compiled all data from the replicates into the CDF and report the error as the precision of the fit. Dwell times of both utrnWT ‘near’ cofilin was slightly longer lived than those far from a cofilin binding event (τ_1 utrnWTnear_ = 0.67 ± 0.04 sec, τ_1 utrnWTfar_ = 0.56 ± 0.01 sec). In contrast, the presence of cofilin had a much stronger effect on utrnLAM with events near cofilin being significantly longer (τ_1 utrnLAMnear_ =1.25 ± 0.05, τ_1 utrnLAMfar_ = 0.81 ± 0.05). The τ_1_ dwell times for utrnΔN ‘near’ cofilin were more similar to those ‘far’ from cofilin (τ_1 utrnΔNnear_ = 0.42 ± 0.03 sec, τ_1 utrnΔNfar_ = 0.54 ± 0.01 sec) (Fig 4D).

**Figure 4:**
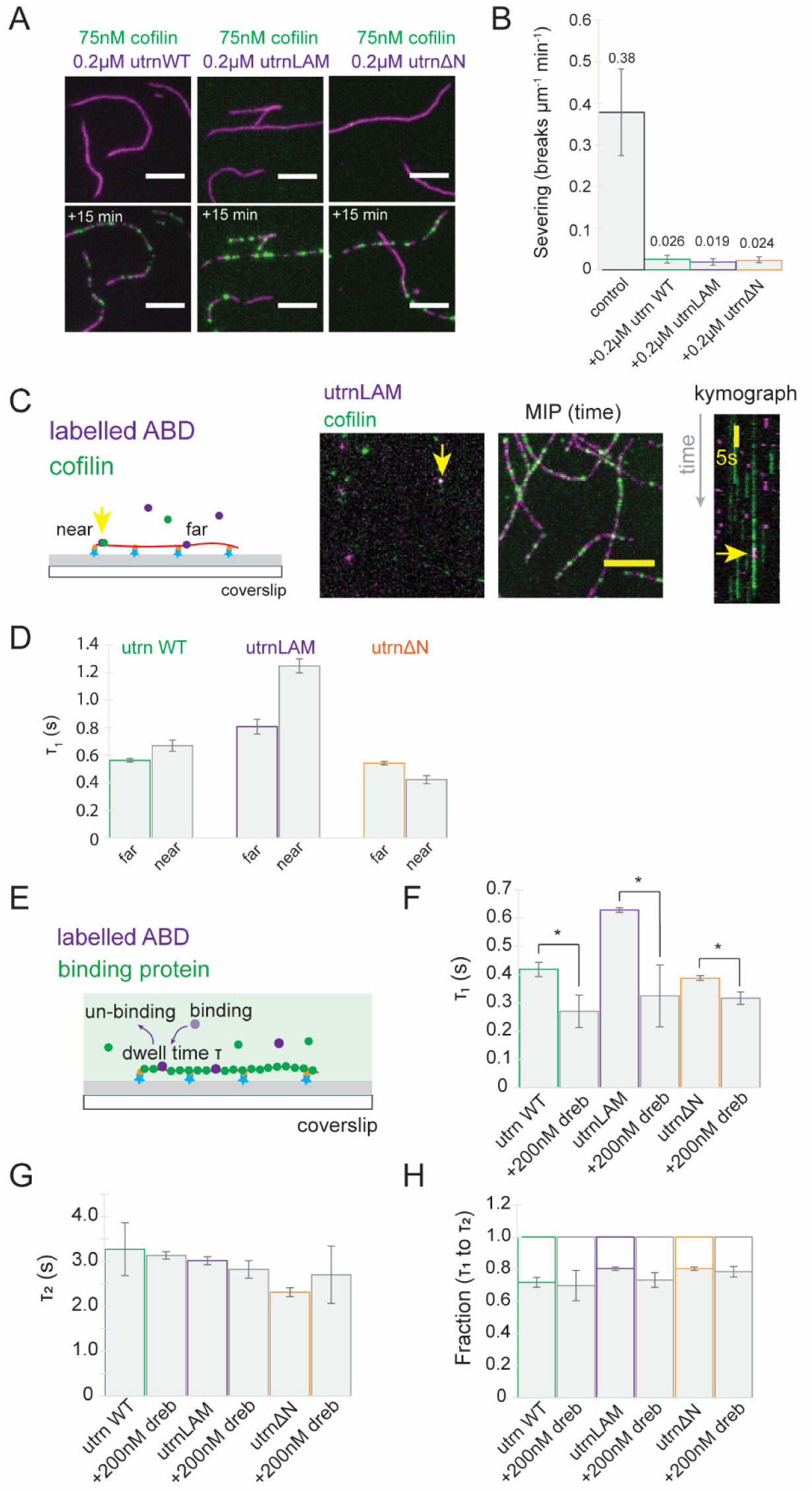
Filament binding by both cofilin and drebrin alters utrophin ABD mutant dwell times. (A) Actin filament severing by cofilin in the presence of 0.2μM of the different utrophin mutants. (B) Quantification of actin filament severing rate. (C) Single molecule colocalization and kinetic measurements at low concentrations of cofilin (10nM) analyzed both near and far from a cofilin cluster. (D) The first timescale from the fit to the CDF for molecules near (<30nm) and far (>30nm) from cofilin for utrn WT, utrnLAM and utrnΔN. (E) Single molecule binding kinetics in the presence of high filament labelling with the actin binding protein drebrin. Comparisons of the first timescale (F) second timescale (G) and relative amplitudes (H) from a double exponential fit to the cumulative distribution functions for the different constructs in the presence of 200nM drebrin.

While cofilin shortens the helical half pitch on actin, the actin binding protein drebrin extends the helical half pitch of an actin filament to a mean of ∼40nm^32,52,55^. Since drebrin does not sever filaments, we were able to use higher concentrations of drebrin and include all binding dwell times in our analysis (Fig 4E). We tested the effect of 200nM drebrin1-300 and found that drebrin binding reduced the dwell time of both utrnWT and utrnLAM (τ_1 utrnWT+dreb_ = 0.27 ± 0.06 sec, p<0.05, τ_1 utrnLAM+dreb_ = 0.32 ± 0.11 sec, p<0.05) (Fig 4F-H). In addition, drebrin binding also had a smaller but significant effect on the binding lifetime of utrnΔN (τ_1 utrnΔN+dreb_ = 0.32 ± 0.02 sec, p<0.05 Fig 4F-H, Fig S4 G-I). Taken together these results show that structural changes induced by actin binding proteins, specifically under-twisting and over-twisting of f-actin, can have an allosteric effect on the kinetic properties of actin binding domains.

### Myosin activity changes the localization and dwell time of utrophin ABD mutants

While cofilin locally remodels actin filaments near the leading edge of migrating cells, myosin generates contractile forces needed for cell migration at the rear of migrating cells^56,57^. Motivated by our observation of differential front-back localization of the utrophin ABD mutants (Fig 1), we tested if myosin activity influenced the binding of purified forms of the utrophin mutants *in vitro*. First, we generated contractile actin networks in vitro using myosin II filaments and α-actinin (Fig 5A). In comparison to a control network (Fig 5B), we found that there were only subtle differences in the localization of utrnWT and utrnLAM. In contrast, utrnΔN was more enriched in actomyosin clusters that utrnWT, while utrnWT and utrnSF displayed the most dramatic differences in localization in actin networks.

**Figure 5:**
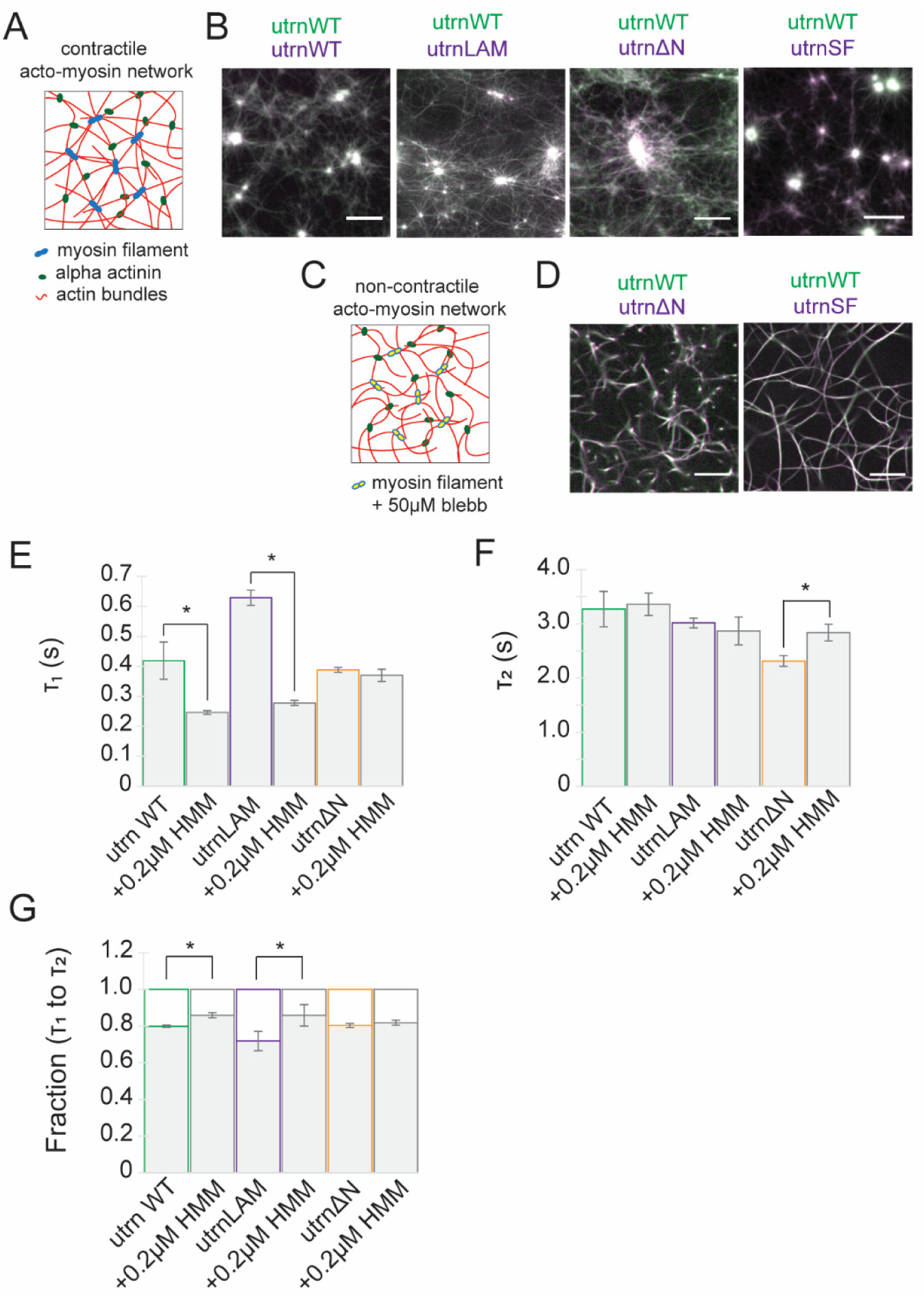
Myosin activity changes localization and dwell time of utrophin ABD mutants. (A) Generation of contractile acto-myosin networks. Localization of (B) utrnWT, utrnLAM, utrnΔN and utrnSF shown in (magenta) relative to utrnWT (green) in actin networks. (C) Generation of non-contractile actin networks through the addition of 50μM blebbistatin. (D) Localization of utrnΔN (magenta) and utrnSF (magenta) relative to utrnWT (green) in a non-contractile gel. Comparisons of the first timescale (E), second timescale (F), and relative amplitudes (G) from a double exponential fit to the cumulative distribution functions for the different constructs in the presence of 200nM HMM.

Interestingly, no significant differences in localization could be observed in gels where contractility was inhibited by blebbistatin (Fig 5C, 5D), suggesting that active myosin was required to cause changes in localization. To investigate the role of myosin activity further, we measured the single molecule dwell times of utrophin ABD mutants in the presence of the myosin fragment Heavy Meromyosin (HMM). We found that utrnWT and utrnLAM displayed a reduced dwell time in the presence of HMM (τ_1 utrnWT+HMM_ = 0.24 ± 0.01 sec, p<0.05, τ_1 utrnLAM+HMM_ = 0.28 ± 0.01 sec, p<0.05, Fig 5E-G, Fig S4 J-L). However, HMM binding had no effect on the binding lifetime of utrnΔN (τ_1 utrnΔN+HMM_ = 0.37 ± 0.02 sec p=0.25). Taken together, these results show that the utrophin mutants have different specificities for actin in the presence of myosin driven contractility.

### Physical confinement of actin alters dwell time of utrophin ABD mutants

Given that conformational changes in f-actin induced by both small molecules and binding proteins impacted the dwell times of utrophin mutants, we next sought to investigate how general this mechanism might be. We tested whether physical constraints on actin filaments to the glass surface influenced the dwell time of single molecules in our assay. To investigate this, we built on our combined single molecule dwell time measurements (kinetics) with sub-pixel localization measurements (STORM) used in our cofilin analysis, to generate images of localized binding dwell-times on actin filaments, an approach we refer to as kSTORM (*materials and methods kSTORM*, Fig 6A).

**Figure 6:**
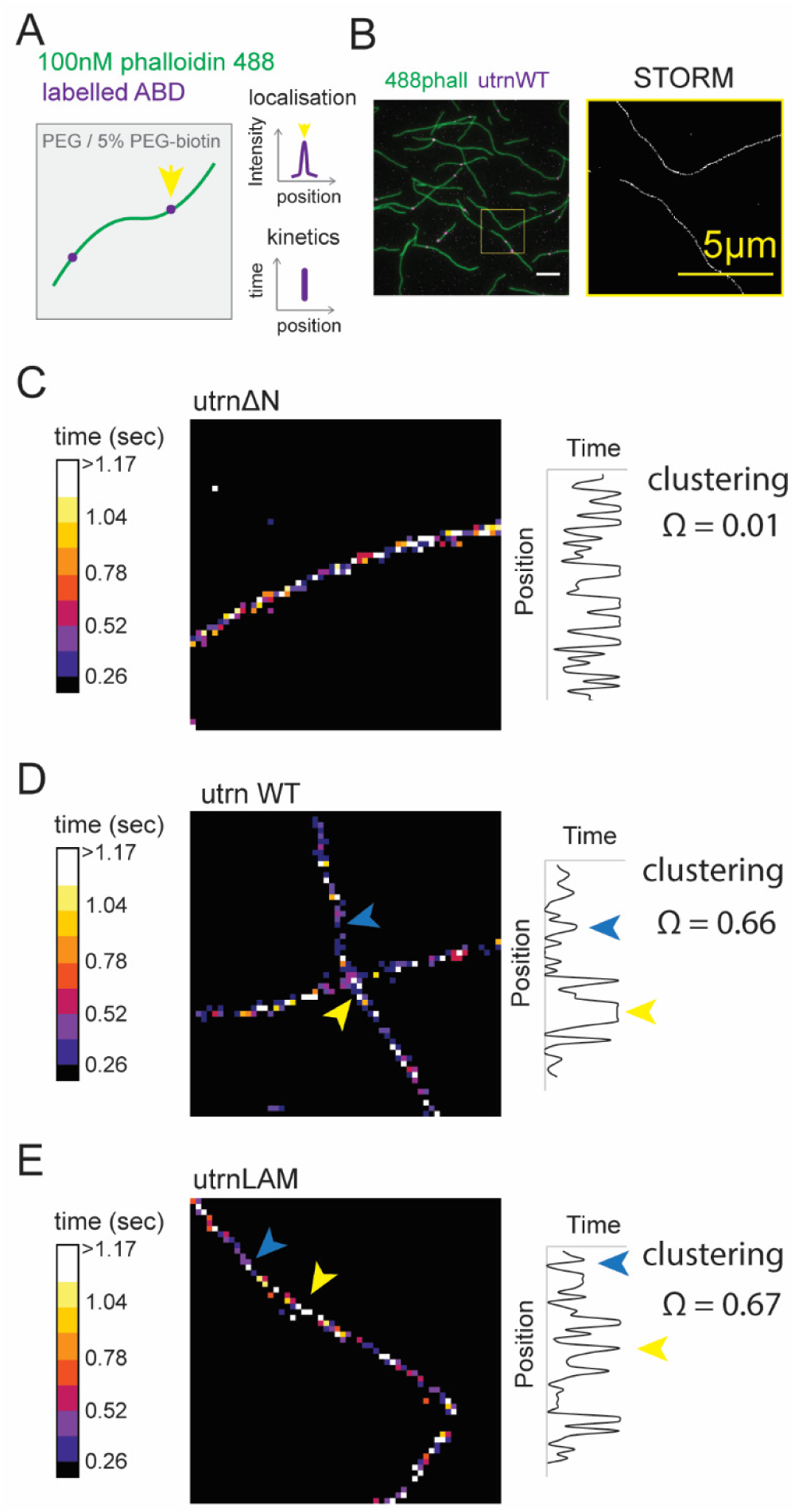
Physical constraints on actin alters dwell time of utrophin ABD mutants. (A) kSTORM combines subpixel localization measurements with kinetics. (B) Example STORM reconstructions of single molecule measurements from utrnWT. (C) kSTORM images for utrnΔN, (D) utrnWT and (E) utrnLAM.

Using kSTORM, we were able to create filament images with a pixel size of 40nm (close to the canonical helical half pitch of f-actin ∼36nm (Fig 6B)) and color-coded the images based on dwell time of the three different utrophin ABD mutants. We observed that the dwell time of utrnΔN was uniformly distributed across filaments (Fig 6C). In contrast, dwell times appeared to cluster for utrnWT (Fig 6D) and utrnLAM (Fig 6E), with short dwell times (<1sec, blue arrowheads) and long dwell times (>1sec, yellow arrowheads) separating into distinct regions. We quantified clustering in these images using a previously reported metric for mixing^58^ and found that utrnWT (Ω_utrnWT_ = 0.66) and utrnLAM (Ω_utrnLAM_ = 0.67) displayed clustering where as utrnΔN did not (Ω_utrnΔN_ = 0.01) (Fig 6).

We hypothesize that as actin filaments are bound to the surface of the flow chamber they adopt a bias for different structural conformations, similar to the dynamic conformational changes in f-actin structure that have been observed using single molecule FRET measurements^24^. This in turn causes clustering of actin binding domain dwell times. These results suggest that mechanical constraint on actin filaments can locally impact the dwell times of different actin binding domain mutants and their segregation into distinct regions.

### Native CH1-CH2 domains display biased localization to different actin structures

Having identified that utrophin ABD mutants localize to different subcellular actin structures and that their binding affinity is altered by changes in actin filament conformation, we wondered if native CH1-CH2 domains displayed similar characteristics. We screened the localization of native CH1-CH2 domains relative to the actin binding domain of utrophin (Fig S5). Native CH1-CH2 domains displayed a range of actin binding affinities, which we assessed from the relative pools of protein on actin and in the cytoplasm in live cells^37^. We also found that several native CH1-CH2 domains displayed enhanced localization to specific actin structures (Fig 7, Fig S5). For example, the actin binding domain of dystonin/BPAG1, a protein that links the actin cytoskeleton to other cytoskeletal networks^59^, was enriched on stress fibers in HeLa cells (Fig 7A, Movie S5). In contrast, the ABD of nesprin II, a protein that links the actin cytoskeleton to the nucleus^60^, was enriched in the lamellipodium in both HeLa cells (Fig 7B, Movie S6) and PLB neutrophils (Fig 7C, Movie S7). These results show that native CH1-CH2 domains, in addition to having different overall affinities, show preferential binding to specific actin structures in cells.

**Figure 7:**
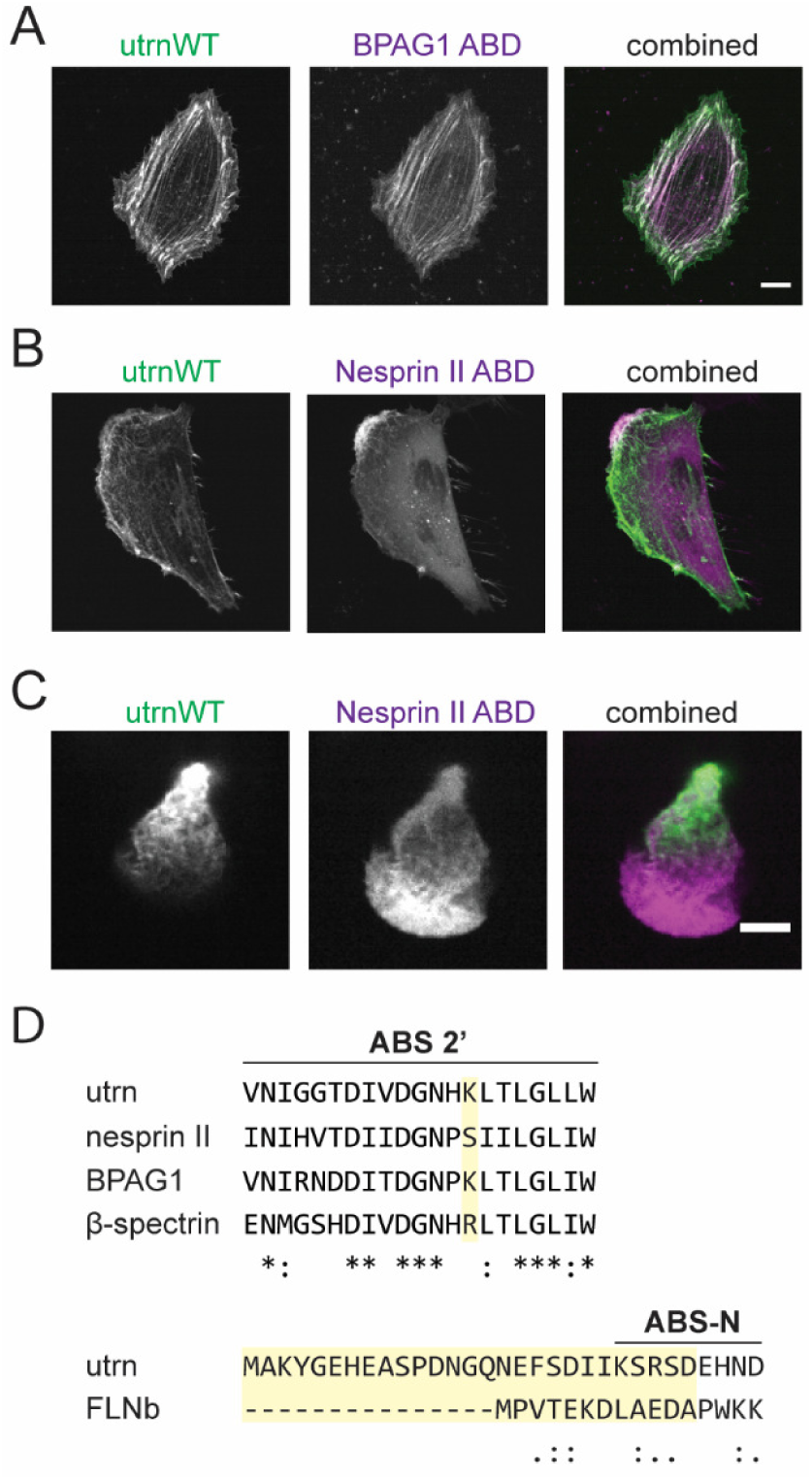
Native CH1-CH2 domains display different sub-cellular localizations. (A) Localization of BPAG1 ABD (magenta) relative to utrnWT (green) in HeLa cells. (B) Localization of Nesprin II ABD (magenta) relative to utrnWT (green). (C) Localization of Nesprin II ABD (magenta) relative to utrnWT (green) in PLB cells. (D) Sequence alignment of native CH1-CH2 domains. Residue K121 for utrnWT highlighted in yellow (top) and the n-terminal region prior to CH1-CH2 and its truncation in yellow (bottom). Identical residues are annotated with ‘*’, strongly conserved with ‘:’, and weakly conserved with ‘.’.

## DISCUSSION

Using a combination of live cell imaging, *in-vitro* characterisation, and single molecule binding measurements, we showed that utrophin ABD mutants have varying binding affinities for different conformational states of f-actin and localization to actin structures in live cells. We found that two constructs, utrnWT and utrnLAM, had different degrees of specificity for structural changes in f-actin, while utrnΔN was largely insensitive to structural changes. These domains responded to biochemical perturbations, regulatory protein binding and interestingly, mechanical constraints on f-actin.

The identification of specificity of actin binding domains to different actin conformations and actin networks has two broad implications for understanding cytoskeletal physiology. Firstly, in addition to generating mutant actin binding domains from utrophin, we tested the localization of native CH1-CH2 domains. Many of these domains displayed differences in binding affinity, characterised by differences in cytoplasmic signal (Fig S5), but several actin-binding domains, including nesprin II CH1-CH2 and BPAG1 CH1-CH2 displayed differences in localization to actin structures. These observations highlight that small differences in sequence between native domains are important for both the affinity and specificity to different actin structures and has broader implications for the activity of full-length actin regulatory proteins. One example of this is indeed nesprin II, which has been shown to localise to the front of the nucleus as cells migrate through small constrictions^60^. This localization was dependent on the presence of the actin-binding domain, suggesting that conformational sensing could help to spatially organise this actin binding protein for its specific function. The broader notion that some actin binding proteins modulate actin filament structure (such as cofilin, formins and myosin), while others can be sensitive to it (CH1-CH2 contain proteins), highlights the role of the actin cytoskeleton as a signalling substrate in its own right, with potential functional significance for a range of biological processes.

Secondly, it is interesting to speculate that CH1-CH2 domains could be used to engineer probes for different structural states of f-actin for use both in vitro and in vivo. In fact, although utrnWT has been commonly used as a marker for f-actin^40^, it has also been reported to localize more preferentially to the trailing edge of migrating cells^61^. C-terminal truncated forms of utrnWT have also been used for labelling of nuclear actin filaments^11^, and direct fusions to GFP via a helical linker have been used in fluorescence polarization studies^62^. We have shown that distinct mechanisms and residues control both affinity^37^ and, in this study, specificity, suggesting that it should be possible to engineer probes for filament conformation with a range of desired properties for live cell and tissue studies^37,63^.

It is important to note that the actin filament conformation-induced differences in ABD localization we report here are distinct from differences in localization that can arise from proteins with different bulk actin-binding affinities. Previous work has shown that high affinity actin-binding proteins such as myosin are depleted from dynamic actin networks due to their slow turnover rate^10,22^. Consistent with this finding, some of the mutants generated in our initial screen had high f-actin binding affinity and displayed differences in localization (*K*_*d*_ */ K*_*d*_ *utrnWT* ∼0.03, utrn Q33A T36A^37^, Fig S5), showing depletion from dynamic actin networks. However, by engineering mutant actin binding domains to have bulk affinities similar to that of utrnWT (Fig 1,2), we are able to specifically identify preferential binding to actin filaments structures independent of bulk affinity differences. Our results suggest that specificity of actin binding proteins to filament conformations could combine with their overall bulk binding affinity to generate a rich landscape of actin binding properties and localizations. This concept could explain why myosin, which binds f-actin with high affinity, also binds cooperatively to actin filaments and displays context-dependent catch bonding behaviour^64,65^, or why highly dynamic Lifeact does not bind to actin decorated with cofilin^13^ or jasplakinolide-stabilized actin^42^, which stresses the filament. In our experiments, the turnover rates of our mutant ABDs (utrnWT, utrnLAM and utrnΔN) are very similar, based on measurements in live cells using FRAP (Fig S6, *Materials and Methods*) and single molecule photoactivation (Fig S7, *Materials and Methods*), even though they display differential localization in cells. In particular, single molecule kinetics in live cells were indistinguishable between utrnWT and utrnLAM when f-actin structures were homogenised with 50µM Y27632 treatment (depolymerises stress fibers, Fig S7). In fact, utrnΔN turned over slightly more slowly than utrnWT, despite its comparative enrichment to more dynamic actin structures (Fig S7E, Fig 1C,D). We also showed that modification of filaments by the actin-stabilizing drugs jasplakinolide and phalloidin have different effects on the binding dwell-time of utrophin ABD mutants in vitro.

How do CH1-CH2 domains sense different conformations of f-actin? Recent evidence has suggested that jasplakinolide preferentially biases one state of f-actin, stabilizing the D-loop from subdomain 2 in a more open configuration^50^, which may partially explain our observations. CH1-CH2 domains have been shown to bind actin by making contacts both on and between actin subunits within the same protofilament^41,42^ (n and n+2). The n-terminal flanking region contacts the n-terminal actin subunit, ABS2 binds within the cleft between the two subunits (where the subdomain 2 D-loop from subunit n, contacts subdomain 1 from subunit n+2) and ABS2’ contacts subunit n+2. Making several contacts on and between neighbouring f-actin subunits could explain why utrophin’s ABD is sensitive to filament level structural changes induced by these small molecule agents. Indeed, the K121A mutation in utrophin corresponds to a key interaction with subdomain 2 and may explain why mutating this residue changes the specificity to jasplakinolide-stabilized actin. Sequence alignment of native CH1-CH2 domains revealed that this residue is well conserved between domains, though some differences do exist (Fig 7D). Interestingly, K121 is changed to serine in nesprin II which also enriched to lamellipodial actin in a similar fashion to utrnLAM (Q33A T36A K121A). In our previous work we have shown that ABS-N is important for localization of CH1-CH2 domains. In particular, Filamin B which has a short n-terminal flanking region displayed a similar localization pattern at focal adhesions to utrnΔN^37^ (Fig 7D). Here, we extend this observation by showing that this region also appears to have a crucial role in actin filament conformational sensing. In all of the conditions tested utrnΔN showed little to no difference in binding dwell time, suggesting it is largely insensitive to differences in f-actin conformation.

In addition to actin drugs, we show that actin binding proteins that modify actin filament conformations such as drebrin, myosin, and cofilin, impact the binding affinity of the different utrophin mutants. Indeed, treatment of live cells expressing utrnWT and utrnLAM with blebbistatin, that inhibits contractility, or calyculin A, that stimulates contractility changed the localization of the different mutants, linking some of our in vitro measurements with those at the cellular level (Fig S8). While in vitro off rate measurements provide a precise measurement of binding rates in a controlled environment, further work will be needed to dissect the contribution of different mechanisms that could influence actin filament conformation in live cells. For example, the role of different actin isoforms was not assessed here.

Surprisingly, protein-induced conformational changes are not required to alter the binding kinetics of our utrn ABD mutants. Indeed, we use kSTORM to make the observation that dwell times on actin filaments physically confined to a glass surface show spatial nonuniformity and clustering for utrnWT, utrnLAM but not for utrnΔN. Our data suggests that filament structural conformations can therefore be biased by physical forces exerted on them by binding of proteins and drugs, as well as direct physical constraints. Actin binding proteins that change the helical pitch of actin, including cofilin, drebrin and formins, exert a torque on the filament^20,36^. This is also the case for myosin II, which steps at a distance shorter than the helical half pitch of an actin filament, causing filaments gliding on a myosin-coated surface to spiral^66,67^. Since actin filaments are inherently helical in nature, torsion and bending are believed to be coupled to twisting^68^ and could arise as filaments are tethered to the glass surface, in a similar fashion to the dynamic conformational changes in actin filaments shown by single molecule FRET^24^. While the mutagenesis study performed here highlights significant new functional roles for different residues on CH1-CH2 domains, further structural work will be needed to identify the binding mechanisms in more detail and how these different regions combine with overall bulk affinity to give rise to unique actin binding properties.

## ACKNOWLEGEMENTS

The authors would like to thank Dyche Mullins and members of the Fletcher lab for helpful discussions. This work was supported by grants from NIH (D.A.F). A.R.H. was in receipt of an EMBO long-term fellowship 1075–2013 and HFSP fellowship LT000712/2014. B.B. was supported by the Ruth L. Kirschstein NRSA fellowship from the NIH (1F32GM115091). P.J was supported by an NSF Fellowship and the Berkeley Fellowship for Graduate Studies. A.B. was supported by the Miller Institute for Basic Research at UC Berkeley as a Miller Visiting Professor. We would like to thank Holly Aaron and Feather Ives at the Molecular Imaging Center core facility at UC Berkeley (FRAP experiments were performed at the Molecular Imaging Center using equipment supported by the Helen Wills Neuroscience Institute). We would also like to acknowledge the help of the Marqusee Lab with the CD and melting temperature experiments. D.A.F. is a Chan Zuckerberg Biohub Investigator.

## MATERIALS AND METHODS

### Cell Culture

HeLa and HEK293 cells were cultured at 37°C in an atmosphere of 5% CO_2_ in air in DMEM (Gibco, #10566024) supplemented with 10% FBS (Gibco, #16140071) and 1% Penicillin-Streptomycin (Gibco, #15140122). Adherent cells were passaged at a 1:5 dilution using 0.05% trypsin EDTA (Gibco #25200056). PLB cells were a kind gift from Dr. Sean Collins (UC Davis). PLB cells were cultured in RPMI (Gibco, #11875093) containing 10% FBS and 1% Penicillin-Streptomycin and differentiated into neutrophil like cells by adding 1.5% DMSO for 5-6 days.

### Generation of constructs

To visualize the relative localization of fluorescent fusions to actin binding domains, we generated both bi-cistronic expression plasmids for transient transfection and two separate lentiviral plasmids for creating double expression stable cell lines, as described previously^37^. Mutations to the actin binding domain of utrophin were introduced by PCR. Briefly, two sets of primers containing the point mutation were used to amplify two separate segments of mCherry-utrn ABD (or in some cases EGFP-utrn ABD or RubyII-utrn ABD) which were then re-assembled using Gibson assembly. Transient transfections were performed using effectene (Qiagen, #301425), following the manufacturer’s protocol and imaged 24 hours after transfection. For generating stable cell lines GFP-utrn ABD and the construct of interest fused to mCherry were cloned into Lentiviral plasmid pHR. Lentiviruses were then generated by transfecting the plasmids into HEK293 cells for viral packaging. Lentiviral supernatants were collected 48hrs after infection, filtered using a 0.4um filter and used directly to infect the target cell line in a 1:1 ratio with normal culture media. PLB cells were infected by centrifuging cells at 300 rcf for 10 minutes in lentiviral supernatant containing polybrene.

### Engineering utrn ABD affinity and specificity

In our previous work^37^, we have shown that two mechanisms control the overall binding affinity of CH1-CH2 domains. Firstly, CH1-CH2 inter-domain interaction govern the ‘openness’ of the two CH domains which relieves a steric clash between CH2 and f-actin. Secondly, CH1-factin interactions govern direct binding to f-actin. Each of these mechanisms can be targeted to titrate the overall actin binding affinity of the domains. The double mutant Q33A T36A perturbs interdomain interactions and makes it easier for the actin binding domain of utrophin to transition to an open bound state on f-actin. These mutations caused an increase in actin binding affinity through changes in both on rate and off rate. By combining inter-CH domain interactions with f-actin binding interactions it is possible to create a range of binding properties tuning affinity to be similar whilst probing different regions on utrn ABD to test specificity to binding to different conformations of f-actin.

### Cellular confocal imaging

Cells expressing fluorescent fusion proteins were imaged using the following excitation and emission: GFP was excited at 488 nm and emission was collected at 525 nm, mCherry was excited at 543 nm, and emission was collected at 617 nm. Live imaging experiments were performed in normal cell culture media using an OKO labs microscope stage enclosure at 37°C in an atmosphere of 5% CO_2_. Cells were imaged on glass bottomed 8 well chambers that had been coated with 10ug/ml fibronectin in PBS for 30minutes. Cells were imaged with a 60x oil immersion objective N.A. 1.4.

### Cellular inhibitor treatments

HeLa cells were treated with either 25µM blebbistatin for 30mins or 0.5nM calyculin A for 15mins. Inhibitor treatments were performed in environmental conditions using an OKO Labs heated microscope stage.

### Fluorescence Recovery After Photobleaching (FRAP)

To assess the turnover kinetics and mobility of utrnABD mutants Fluorescence Recovery After Photobleaching experiments were performed. FRAP measurements were performed specifically on stress fibers in HeLa cells. The turnover of different mutants was measured by bleaching a 6pixel diameter spot (∼1µm) using a scanning laser confocal microscope (Zeiss LSM 880 with Airyscan). Fusion constructs to mCherry were used in FRAP experiments. To analyse FRAP data, time lapse stacks were imported into Fiji and bleached regions analysed as ROI. FRAP data were bleaching corrected as previously described^70^ and the initial rate of recovery found from the initial slope of the recovery curve using MatLab.

### Single molecule binding measurements in live cells

To complement the kinetic measurements in live cells using FRAP on stress fibers, we used photoconversion and single molecule binding measurements. Mutants of interested were generated as fusions to mEOS for single molecule photoactivation with TIRF microscopy. Because cells contain a range of different actin structures that could influence the binding kinetics results, we pre-treated cells with 50µM of the ROCK inhibitor Y27632 for 30mins, to depolymerize stress fibers. Single molecules were then activated with a 30ms pulse of 405nm light in TIRF, and then imaged with 561nm excitation at an interval of 50ms. Single molecules were identified and tracked using the TrackNTrace software package^71^. A custom written MatLab routine was then used to post-process the image tracks and calculate binding dwell-times.

### Protein purification and labelling

Actin was purified from rabbit muscle acetone powder (Pel Freez Biologicals, #41995-1) as previously reported^72^, and stored in monomeric form in G-buffer (2mM Tris-Cl pH 8.0, 0.2 mM ATP, 0.5 mM TCEP, 0.1 mM CaCl_2_) at 4°C. Utrophin’s actin binding domain (CH1-CH2) and its associated mutants were expressed recombinantly in E. coli BL21 (DE3) pLysS (Promega, #L1191). Cells were lysed by sonication and HIS tagged protein containing a SUMO solubility tag were purified using affinity chromatography. The solubility tag was cleaved off using TEV protease which was also his tagged, and removed by recirculation over the affinity column. Finally, proteins were purified by size exclusion chromatography. Proteins were stored in 20 mM Tris-Cl pH 7.5, 150 mM KCL, 0.5 mM TCEP and 0.1 mM EDTA and in the presence of 20% glycerol. Utrophin ABD sequences included a KCK linker (GGSGKCKSA) on the C terminus for labelling. Proteins were labelled using either Alexa 488 C_5_ maleimide, Alexa 555 C_2_ maleimide or Alexa 647 C_2_ maleimide (ThermoFisher, #A10254, #A20346, #A20347) as previously described^37^. The minimal actin binding portion of drebrin 1-300 was purified using the same strategy. Acanthamoeba α-actinin and Atto-488 cofilin were a kind gift from Peter Bieling (Max Plank Institute of Molecular Physiology, Dortmund).

### Surface functionalization and flow well assembly

Crosslinked network and single filament assays were performed in a flow well configuration consisting of a functionalized coverslip and passivated counter-surface assembled using Tesa double sided tape. Glass slides (VWR, #48300-047) were plasma cleaned then passivated using PLL-PEG (g = 3.5). 22×22mm coverslips (Zeiss, #474030-9020-000) were passivated using PEG-silane chemistry^73^. Firstly, glass coverslips were cleaned with 3N NaOH, rinsed in miliQ water, piranha cleaned, rinsed and dried, and then incubated with GOTPS for 1 hour at 75°C. After silanizaion, the coverslips were rinsed in anhydrous acetone and dried. PEG was coupled to the silanized surface by preparing a PEG saturated acetone solution at 95% hydroxy-amino-PEG (Rapp Polymere, #10 3000-20) and 5% biotinyl-amino-PEG (Rapp Polymere, #13 3000-25-20) which was incubated for a minimum of 4 hours at 50°C. PEG passivated coverslips were then rinsed in miliQ and stored at room temperature and used within 1 month.

### Total Internal Reflection Fluorescence Microscopy (TIRF)

TIRF microscopy was used for measuring single molecule binding kinetics in cells and *in vitro*. The imaging system consisted of a Nikon TIRF inverted scope (Nikon Eclipse Ti, 488/560/642nm OPSL lasers) with perfect focus, a 100x N.A. 1.4 APO TIRF oil objective, and an EMCCD camera (Andor iXon Ultra).

### In vitro single molecule binding kinetics assay

To evaluate the binding properties of different utrophin ABD mutants, single molecule binding kinetics were measured. Actin filaments were polymerized at a final concentration of 5 µM at room temperature. To immobilize actin filaments to the surface of the flow chamber, flow wells were first incubated with 10µg/mL streptavidin (Sigma #S0677) for 1 minute, washed with f-buffer and then incubated with 1µM biotin phalloidin, (ThermoFisher #B7474) for 1 minute. Actin filaments were then diluted 50x in fbuffer and immediately introduced into the flow well and allowed to attach for 5 minutes. Remaining filaments were washed away with assay buffer (25mM Immidizole, 25mM KCl, 4mM MgCl_2_, 1mM EGTA, 1mM DTT and 10µg/mL Beta Casein (Sigma C6905)). Binding proteins were diluted to a sufficiently low concentration to enable the visualisation of single molecules in TIRF, 0.05-10nM in assay buffer. For single molecule kinetic measurements 600 frames were acquired at an interval of ∼30-130ms depending on the construct.

### Single molecule analysis

Single molecules were identified and tracked using the TrackNTrace software package^71^. A custom written MatLab routine was then used to post-process the particle tracks and calculate binding dwell-times. As a first step, a maximum intensity projection (MIP) through time of the single molecule movies was used to identify the filament backbone (Fig 2A). An image mask was generated from the MIP by thresholding above the background intensity, and made contiguous by image closure. The MIP mask was then used to filter out single binding events in the maximum intensity projection which did not reside within filament backbone (non-specific binding events). Non single molecule binding events were also removed by setting a maximum intensity threshold. Both of these filters excluded a small fraction of total events (Fig 2A, Fig S3). Binding measurements were then calculated from single molecule tracks that occurred within the filament masked regions. To calculate the binding on rate, the length of actin filaments within an image was calculated from the MIP mask by skeletonization of the mask. On rate was then calculated as the total number of events that occurred during the time of the single molecule movie, for a given number of available binding sites, as described previously^48^. For dwell time measurements, the population of recovered single molecule binding events for different actin binding mutants were analysed in two different ways. Firstly, the average dwell time (τ_av_) for the entire population was measured as a metric for bulk binding dwell time. Secondly, the cumulative distribution function of binding dwell times was calculated and fitted with a two-timescale binding model. 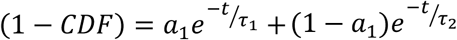. Pearson’s correlation coefficient was used to compare single versus two-timescale models.

### kSTORM analysis

To evaluate local differences in dwell time that occur along single filaments, we combined STORM resolution with dwell time measurements. A similar approach to the single molecule binding kinetics measurement was used with some additional modifications. Due to the number of frames required to construct a STORM image, an additional channel was used to correct for x-y drift during image acquisition. We used 488phalloidin for this reference channel and registered drift in the image stack using this channel. A phalloidin image was captured every 600 frames for a total of 12,000 single molecule images. TrackNTrace was used to fit the single molecule locations using the inbuilt wavelet filtering algorithm. Dwell times were extracted as for single molecule kinetic measurements and reconstructed back into an image using the super-resolution localizations. To compare clustering in the different kSTORM images the mixing parameter Ω that has been previously used to analyse sequence data^58^.

### Actin network assays

To generate actin networks, all reagents and binding proteins, excluding g-acitn, were added in presence of AB buffer (25mM Immidizole, 25mM KCl, 4mM MgCl_2_, 1mM EGTA, 1mM DTT). Actin was then added to initialize network formation. Actin and binding proteins were used at final concentrations of: 12µM actin, 200nM of each fluorescently labelled actin binding domain, 2.5µM α-actinin and 500nM myosin II. Before actin polymerisation, samples were incubated for 5 minutes to allow for the assembly of myosin filaments and homogenized by pipetting to obtain a near uniform myosin filament size before adding to the imaging chamber. Samples were incubated in the imaging chamber for 3 minutes to generate contractile networks and then images immediately with spinning disc confocal microscopy. To generate non-contractile networks, 50µM blebbistatin was added to the initial mix, which inhibits contractility but not myosin filament assembly.

### Statistics

Error bars represent standard error, unless otherwise specified. Statistical significance was determined by a two-tailed student’s t-test and assumed significant when p<0.05. For single molecule dwell time measurements, individual replicates were considered to be individual imaging chambers (consisting of >1000 binding events) imaged on different days. A minimum of 3 replicates was measured for each condition. For single molecule measurements in the presence of cofilin, coincident events were rare (<1% of total binding events). To compare cofilin measurements the CDF was assembled by combining all of the different replicates. In these measurements error bars represent the 95% confidence interval of the CDF fit.

## SUPPLEMENTARY FIGURES

**Figure S1:**
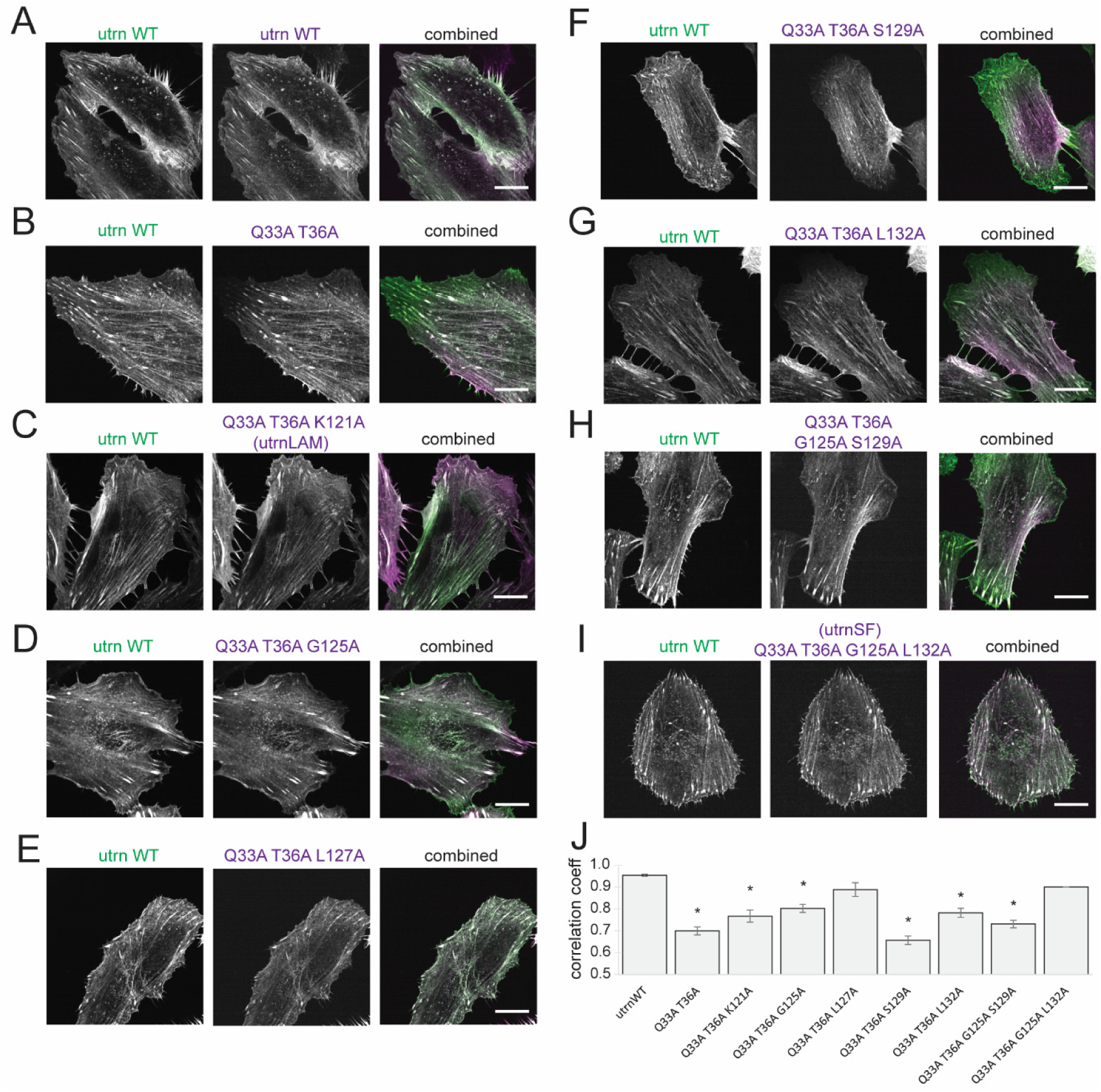
Localization screen of utrn CH1 mutants. (A-I) Localization of alanine mutations to different residues on utrn CH1 shown in HeLa cells. utrnWT is shown in green and the mutant in magenta. (J) Quantification of localization by Pearson’s correlation coefficient for the two color channels.

**Figure S2:**
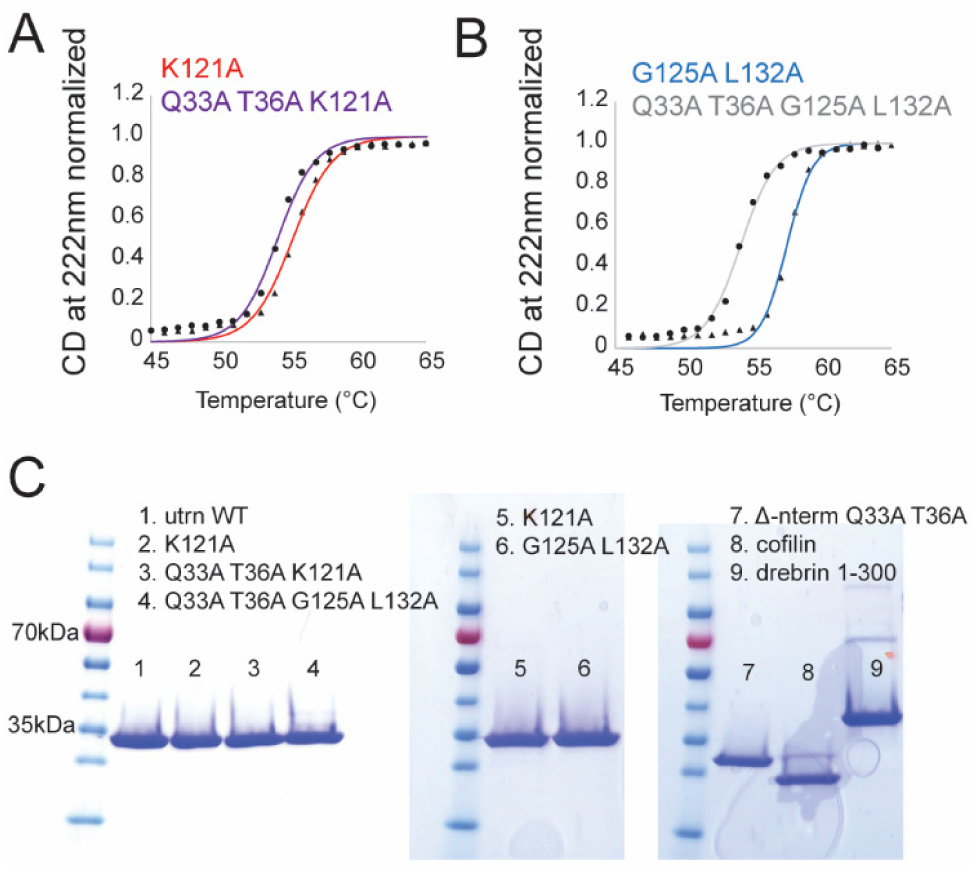
In vitro characterization of utrn CH1-CH2 mutants. (A) Melting temperature curves for K121A (red) and Q33A T36A K121A (purple). (B) Melting temperature curves for G125A L132A (blue) and Q33A T36A G125A L132A (grey). (C) SDS page gels for the purified proteins.

**Figure S3:**
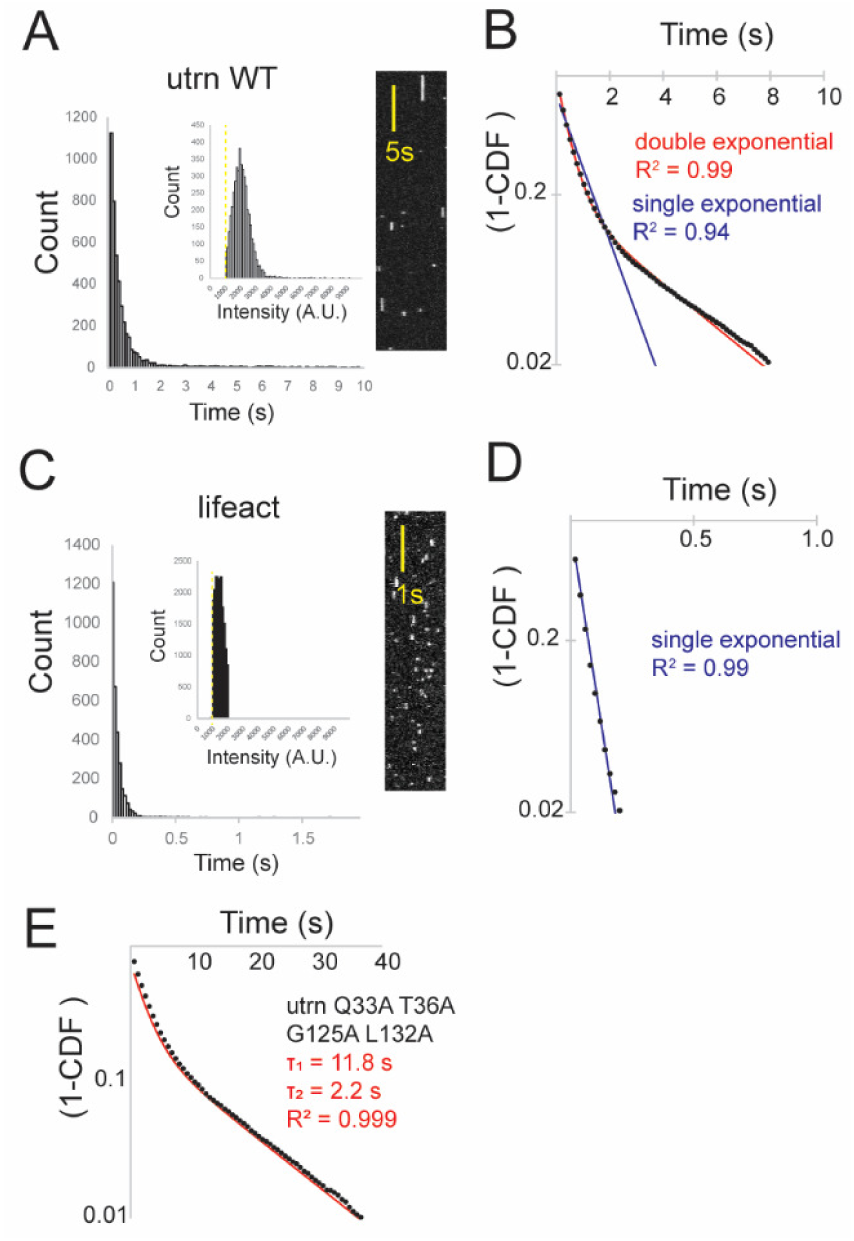
Calibration of single molecule binding kinetics. (A) Dwell time histogram, intensity histogram and kymograph for utrnWT. (B) Cumulative distribution function for utrnWT fit with either a single (blue) or double exponential decay (red). (C) Dwell time histogram, intensity histogram and kymograph for lifeact. (D) Cumulative distribution function for lifeact, fit with a single exponential. (E) Binding timescale as a function of concentration of protein added. (F) Cumulative distribution function for utrn Q33A T36A G125A L132A fit with a double exponential.

**Figure S4:**
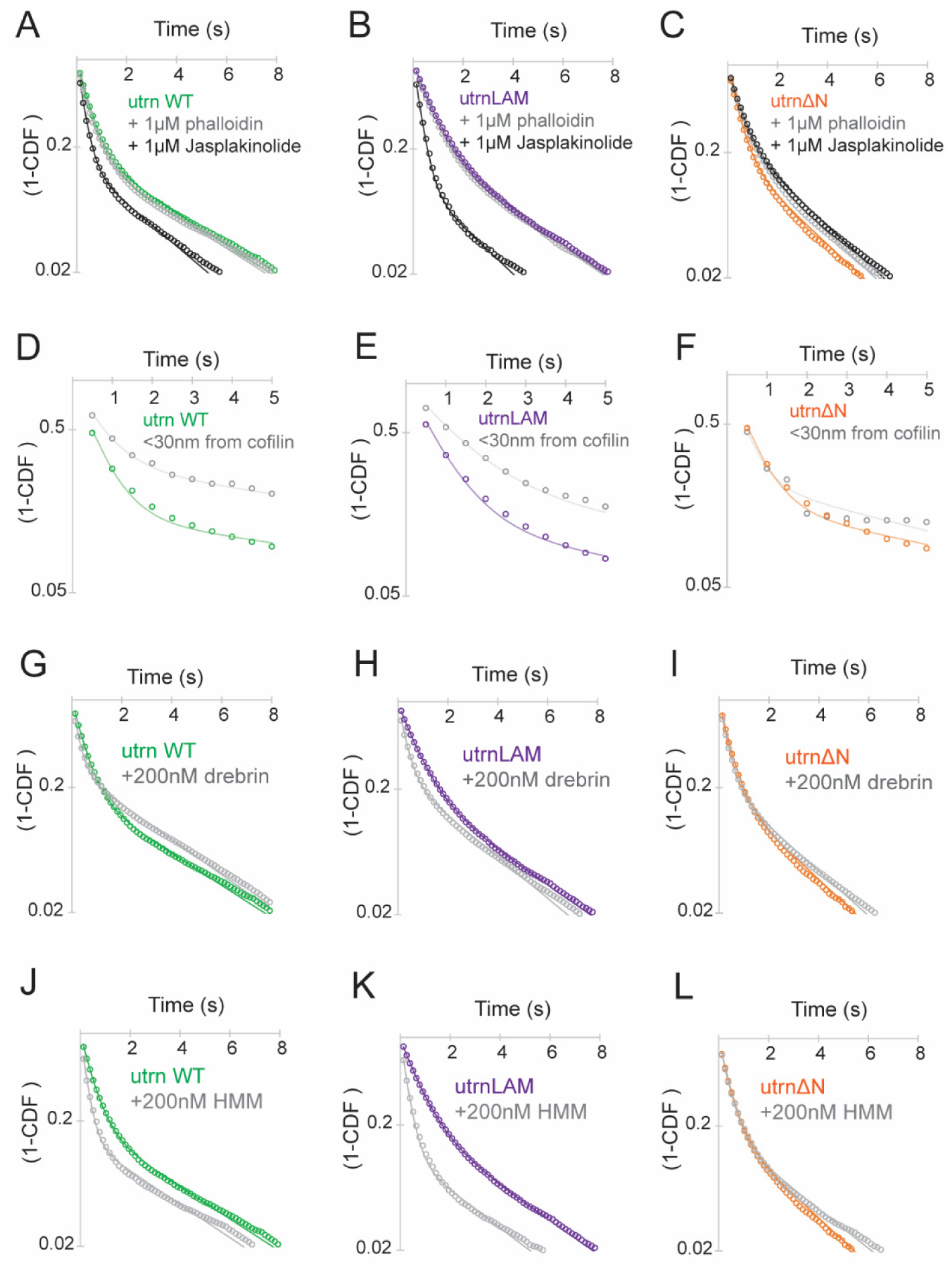
Cumulative distribution functions for different constructs and conditions. (A-C) In the presence of either 1µM phalloidin or 1µM jasplakinolide. (D-F) near and far from cofilin. (G-I) In the presence of 200nM drebrin. (J-L) In the presence of 200nM HMM.

**Figure S5:**
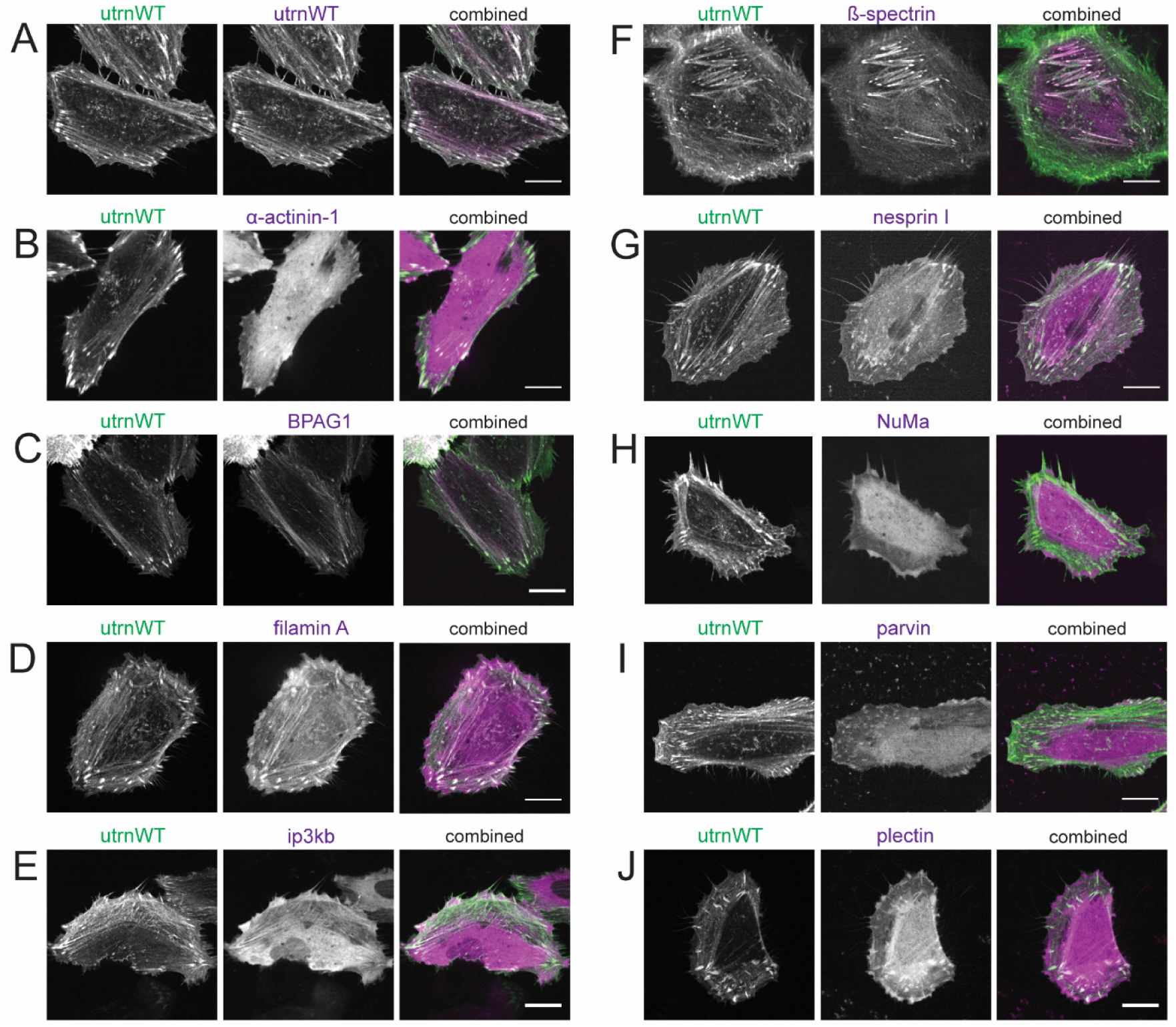
Localization of native CH1-CH2 domains. Localization of native CH1-CH2 domains shown in magenta relative to utrnWT shown in green. (A) utrnWT, (B) α-actinin 1, (C) BPAG1, (D) Filamin A, (E) ip3kb, (F) β-spectrin, (G) nesprin I, (H) NuMa, (I) parvin, (J) plectin.

**Figure S6:**
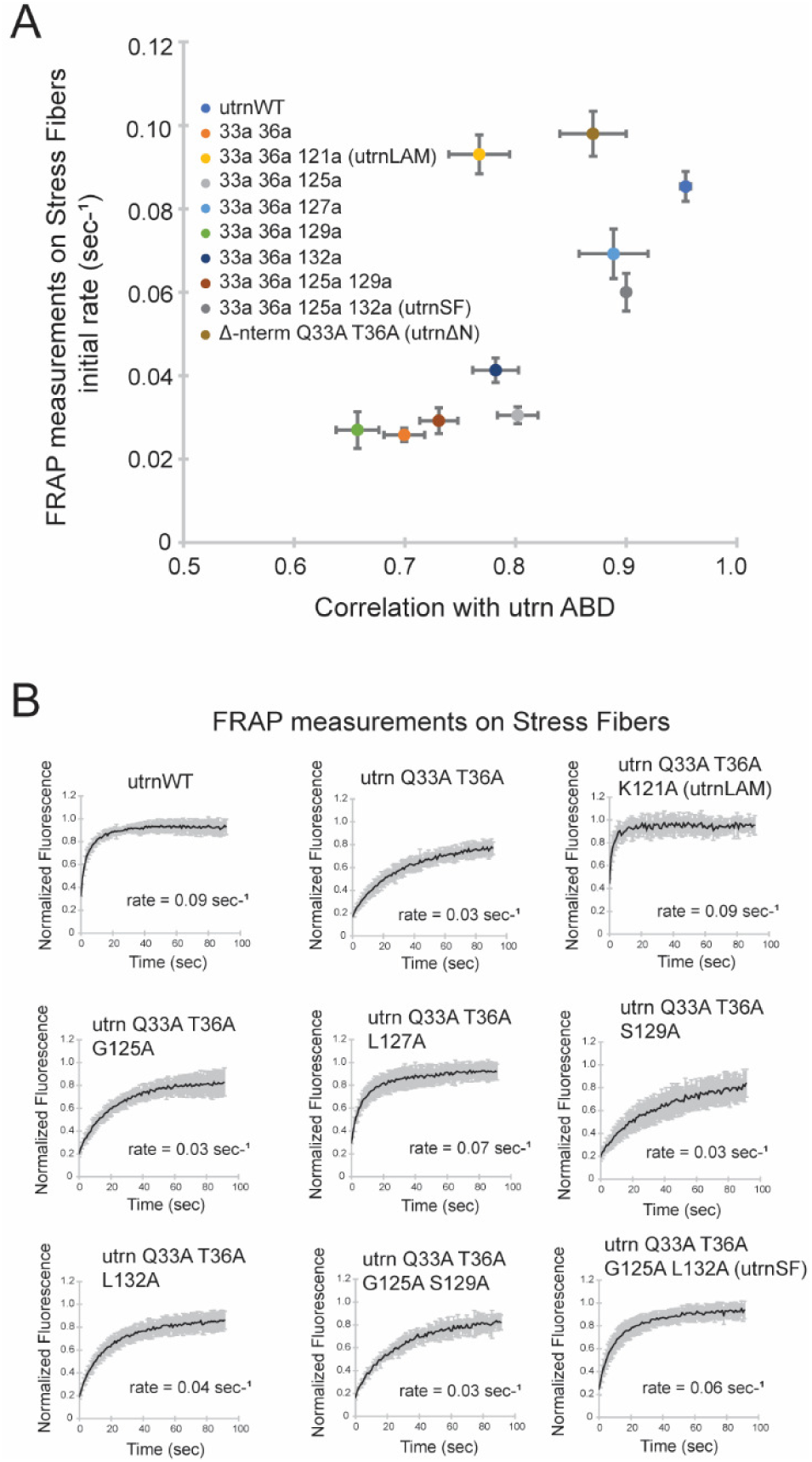
Comparison of localization and recovery rate on stress fibers for utrn CH1-CH2 mutants. (A) Comparison of Pearson’s correlation coefficient and the initial recovery rate of different utrn CH1-CH2 mutants measured via Fluorescence Recovery After Photobleaching (FRAP) on stress fibers in HeLa cells. utrnΔN recovery and localization values from Harris et al^37^. (B) FRAP recovery curves from measurements on stress fibers.

**Figure S7:**
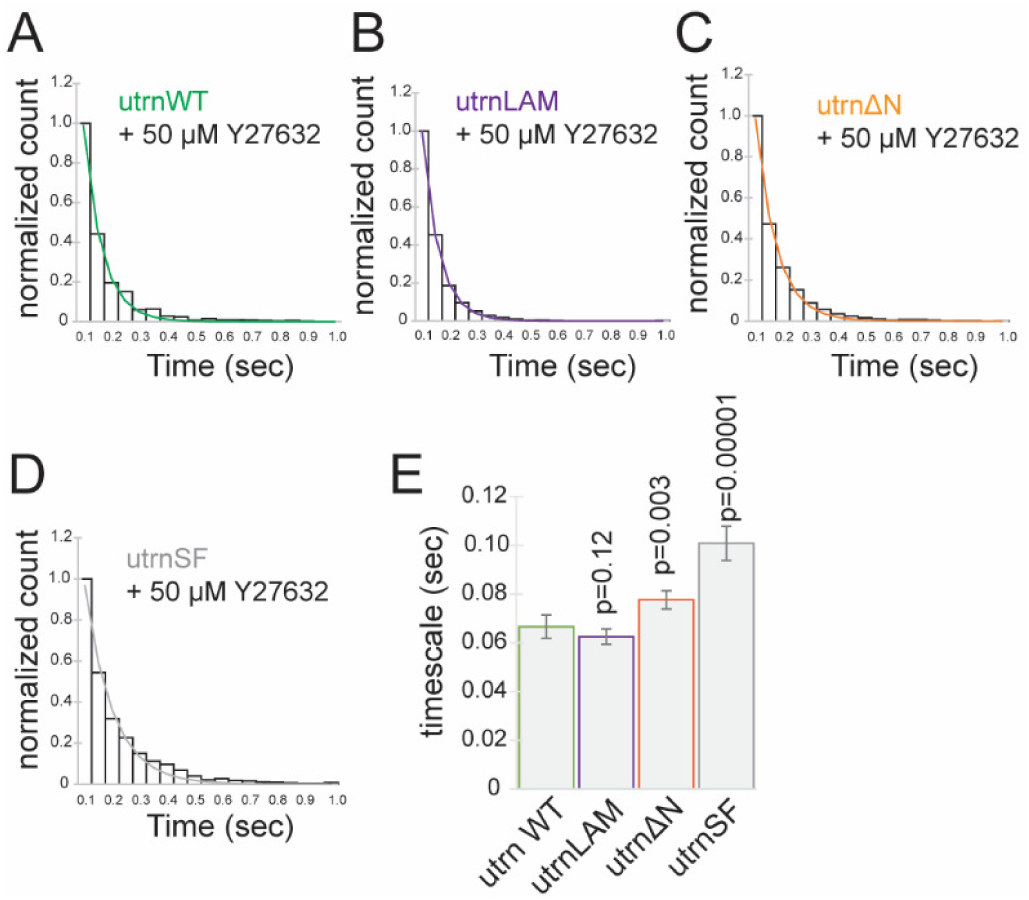
Kinetic measurements of utrn CH1-CH2 mutants in live cells. Single molecule dwell time histogram measured by photoactivation of utrophin mutant fusions to EOS in HeLa cells treated with 50µM Y27632 for (A) utrnWT, (B) utrnLAM, (C) utrnΔN and (D) utrnSF. (E) measurements of binding timescales for the different constructs.

**Figure S8:**
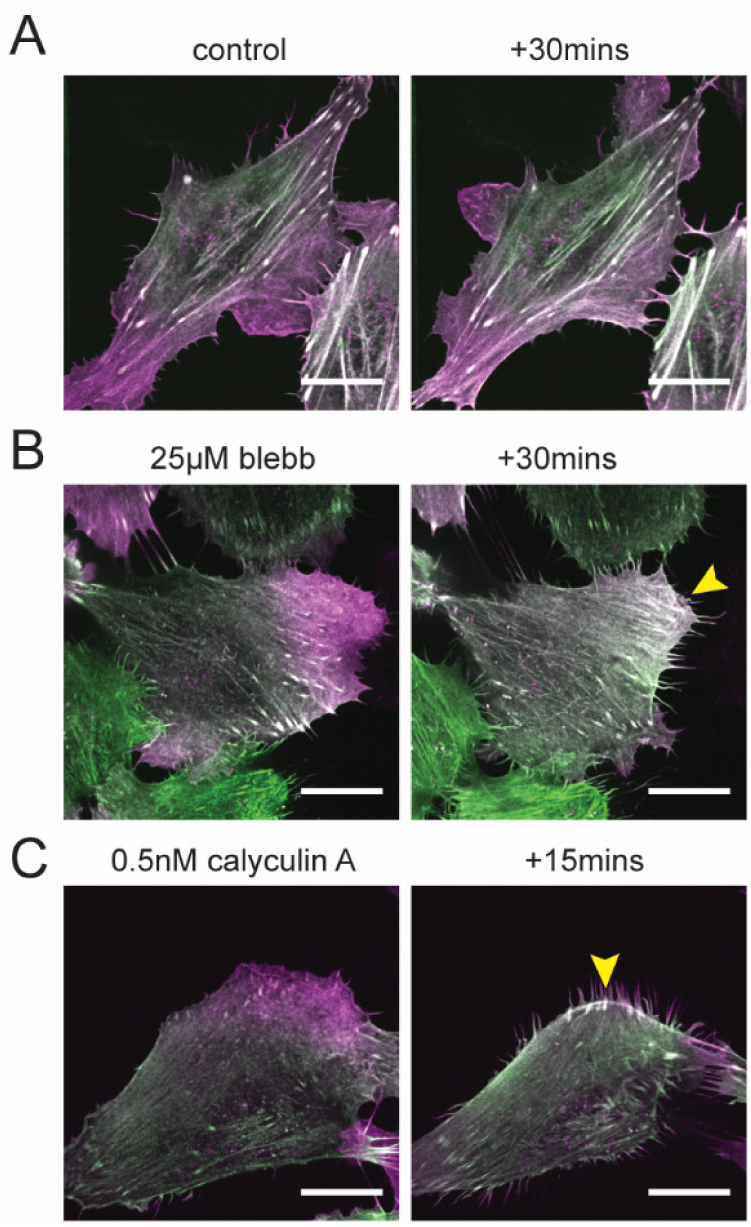
Cellular inhibitor measurements. In all images utrnWT is shown in green and utrnLAM is shown in magenta. (A) Control cells. (B) Cells treated with 25µM blebbistatin for 30 minutes. (C) Cells treated with 0.5nM calyculin A for 15 minutes. Scale bar in all images is 20µm.

## SUPPLEMENTARY MOVIES

**Movie S1:**
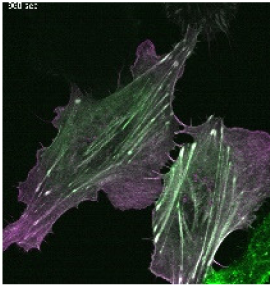
HeLa cell, utrnWT (green) – utrn Q33A T36A K121A (magenta)

**Movie S2:**
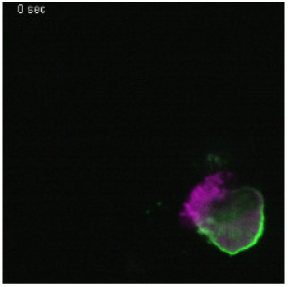
PLB cell, utrnWT (green) – utrn Q33A T36A K121A (magenta)

**Movie S3:**
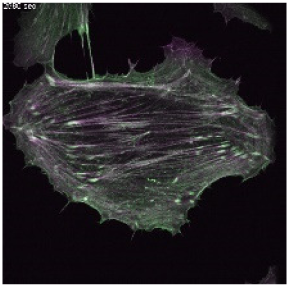
HeLa cell, utrnWT (green) – Δ-nterm utrn Q33A T36A (magenta)

**Movie S4:**
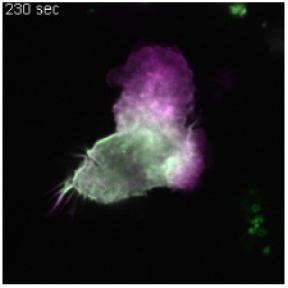
PLB cell, utrnWT (green) – Δ-nterm utrn Q33A T36A (magenta)

**Movie S5:**
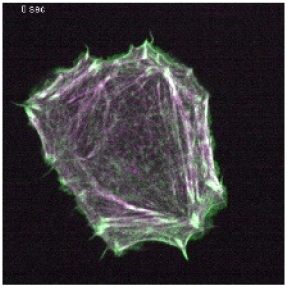
HeLa cell, utrnWT (green) – BPAG1 ABD (magenta)

**Movie S6:**
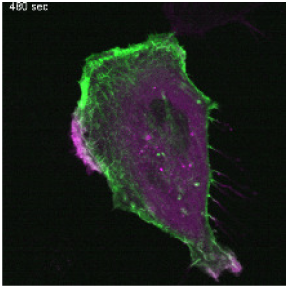
HeLa cell, utrnWT (green) – Nesprin II ABD (magenta)

**Movie S7:**
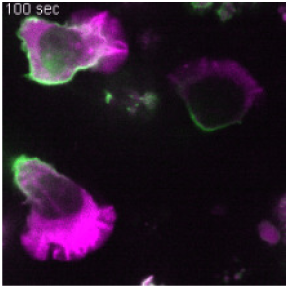
PLB cell, utrnWT (green) – Nesprin II ABD (magenta)

## REFERENCES

1. Michelot, A. & Drubin, D. G. Building distinct actin filament networks in a common cytoplasm. Curr. Biol. 21, R560–R569 (2011).

2. Harris, A. R., Jreij, P. & Fletcher, D. A. Mechanotransduction by the Actin Cytoskeleton: Converting Mechanical Stimuli into Biochemical Signals. Annu. Rev. Biophys. 47, 617–631 (2018).

3. Svitkina, T. M. et al. Mechanism of filopodia initiation by reorganization of a dendritic network. J Cell Biol 160, 409–421 (2003).

4. Fritzsche, M. et al. Self-organizing actin patterns shape membrane architecture but not cell mechanics. Nat. Commun. 8, 1–14 (2017).

5. Luxenburg, C. et al. The architecture of the adhesive apparatus of cultured osteoclasts: from podosome formation to sealing zone assembly. PloS One 2, e179 (2007).

6. Young, M. E., Cooper, J. A. & Bridgman, P. C. Yeast actin patches are networks of branched actin filaments. J. Cell Biol. 166, 629–635 (2004).

7. Riedl, J. et al. Lifeact: a versatile marker to visualize F-actin. Nat. Methods 5, 605–607 (2008).

8. Schell, M. J., Erneux, C. & Irvine, R. F. Inositol 1, 4, 5-trisphosphate 3-kinase A associates with F-actin and dendritic spines via its N terminus. J. Biol. Chem. 276, 37537–37546 (2001).

9. Lopata, A. et al. Affimer proteins for F-actin: novel affinity reagents that label Factin in live and fixed cells. Sci. Rep. 8, 6572 (2018).

10. Yamashiro, S. et al. Convection-induced biased distribution of actin probes in live cells. Biophys. J. 116, 142–150 (2019).

11. Belin, B. J., Goins, L. M. & Mullins, R. D. Comparative analysis of tools for live cell imaging of actin network architecture. BioArchitecture 4, 189–202 (2014).

12. Melak, M., Plessner, M. & Grosse, R. Actin visualization at a glance. J Cell Sci 130, 525–530 (2017).

13. Munsie, L. N., Caron, N., Desmond, C. R. & Truant, R. Lifeact cannot visualize some forms of stress-induced twisted F-actin. Nat. Methods 6, 317–317 (2009).

14. Washington, R. W. & Knecht, D. A. Actin binding domains direct actin-binding proteins to different cytoskeletal locations. BMC Cell Biol. 9, 10 (2008).

15. Lemieux, M. G. et al. Visualization of the actin cytoskeleton: different F-actin-binding probes tell different stories. Cytoskeleton 71, 157–169 (2014).

16. Tsujioka, M. et al. Talin couples the actomyosin cortex to the plasma membrane during rear retraction and cytokinesis. Proc. Natl. Acad. Sci. 109, 12992–12997 (2012).

17. Shibata, K., Nagasaki, A., Adachi, H. & Uyeda, T. Q. Actin binding domain of filamin distinguishes posterior from anterior actin filaments in migrating Dictyostelium cells. Biophys. Physicobiology 13, 321–331 (2016).

18. Christensen, J. R. et al. Competition between Tropomyosin, Fimbrin, and ADF/Cofilin drives their sorting to distinct actin filament networks. Elife 6, e23152 (2017).

19. Gateva, G. et al. Tropomyosin isoforms specify functionally distinct actin filament populations in vitro. Curr. Biol. 27, 705–713 (2017).

20. Mizuno, H., Tanaka, K., Yamashiro, S., Narita, A. & Watanabe, N. Helical rotation of the diaphanous-related formin mDia1 generates actin filaments resistant to cofilin. Proc. Natl. Acad. Sci. 201803415 (2018).

21. Risca, V. I. et al. Actin filament curvature biases branching direction. Proc. Natl. Acad. Sci. 109, 2913–2918 (2012).

22. Maiuri, P. et al. Actin flows mediate a universal coupling between cell speed and cell persistence. Cell 161, 374–386 (2015).

23. Galkin, V. E., Orlova, A., Schröder, G. F. & Egelman, E. H. Structural polymorphism in F-actin. Nat. Struct. Mol. Biol. 17, 1318–1323 (2010).

24. Kozuka, J., Yokota, H., Arai, Y., Ishii, Y. & Yanagida, T. Dynamic polymorphism of single actin molecules in the actin filament. Nat. Chem. Biol. 2, 83 (2006).

25. Egelman, E. H., Francis, N. & DeRosier, D. J. F-actin is a helix with a random variable twist. Nature 298, 131 (1982).

26. Pfaendtner, J., Branduardi, D., Parrinello, M., Pollard, T. D. & Voth, G. A. Nucleotide-dependent conformational states of actin. Proc. Natl. Acad. Sci. 106, 12723–12728 (2009).

27. Hung, R.-J., Pak, C. W. & Terman, J. R. Direct redox regulation of F-actin assembly and disassembly by Mical. Science 334, 1710–1713 (2011).

28. Jegou, A. & Romet-Lemonne, G. The many implications of actin filament helicity. in Seminars in Cell & Developmental Biology (Elsevier, 2019).

29. Jégou, A. & Romet-Lemonne, G. Single Filaments to Reveal the Multiple Flavors of Actin. Biophys. J. 110, 2138–2146 (2016).

30. Ngo, K. X. et al. Allosteric regulation by cooperative conformational changes of actin filaments drives mutually exclusive binding with cofilin and myosin. Sci. Rep. 6, 35449 (2016).

31. Umeki, N., Hirose, K. & Uyeda, T. Q. Cofilin-induced cooperative conformational changes of actin subunits revealed using cofilin-actin fusion protein. Sci. Rep. 6, (2016).

32. Sharma, S., Grintsevich, E. E., Hsueh, C., Reisler, E. & Gimzewski, J. K. Molecular cooperativity of drebrin1-300 binding and structural remodeling of Factin. Biophys. J. 103, 275–283 (2012).

33. Mei, L., de los Reyes, S. E., Reynolds, M. J., Liu, S. & Alushin, G. M. Molecular mechanism for direct actin force-sensing by α-catenin. bioRxiv (2020).

34. Galkin, V. E., Orlova, A. & Egelman, E. H. Actin filaments as tension sensors. Curr. Biol. 22, R96–R101 (2012).

35. Hayakawa, K., Tatsumi, H. & Sokabe, M. Actin filaments function as a tension sensor by tension-dependent binding of cofilin to the filament. J. Cell Biol. 195, 721–727 (2011).

36. Wioland, H., Jegou, A. & Romet-Lemonne, G. Torsional stress generated by ADF/cofilin on cross-linked actin filaments boosts their severing. bioRxiv 380113 (2018).

37. Harris, A. R. et al. Steric regulation of tandem calponin homology domain actin-binding affinity. Mol. Biol. Cell mbc. E19-06-0317 (2019).

38. Tojkander, S., Gateva, G. & Lappalainen, P. Actin stress fibers–assembly, dynamics and biological roles. J Cell Sci 125, 1855–1864 (2012).

39. Biro, M. et al. Cell cortex composition and homeostasis resolved by integrating proteomics and quantitative imaging. Cytoskeleton 70, 741–754 (2013).

40. Burkel, B. M., Von Dassow, G. & Bement, W. M. Versatile fluorescent probes for actin filaments based on the actin-binding domain of utrophin. Cell Motil. Cytoskeleton 64, 822–832 (2007).

41. Iwamoto, D. V. et al. Structural basis of the filamin A actin-binding domain interaction with F-actin. Nat. Struct. Mol. Biol. 1 (2018).

42. Kumari, A., Kesarwani, S., Javoor, M. G., Vinothkumar, K. R. & Sirajuddin, M. Structural insights into filament recognition by cellular actin markers. bioRxiv 846337 (2019).

43. Galkin, V. E., Orlova, A., Salmazo, A., Djinovic-Carugo, K. & Egelman, E. H. Opening of tandem calponin homology domains regulates their affinity for Factin. Nat. Struct. Mol. Biol. 17, 614–616 (2010).

44. Singh, S. M., Bandi, S. & Mallela, K. M. The N-Terminal flanking region modulates the actin binding affinity of the utrophin tandem calponin-homology domain. Biochemistry 56, 2627–2636 (2017).

45. Avery, A. W. et al. Structural basis for high-affinity actin binding revealed by a β-III-spectrin SCA5 missense mutation. Nat. Commun. 8, 1350 (2017).

46. Jansen, S. et al. Single-molecule imaging of a three-component ordered actin disassembly mechanism. Nat. Commun. 6, 7202 (2015).

47. Bombardier, J. P. et al. Single-molecule visualization of a formin-capping protein ‘decision complex’at the actin filament barbed end. Nat. Commun. 6, 8707 (2015).

48. Hayakawa, K., Sakakibara, S., Sokabe, M. & Tatsumi, H. Single-molecule imaging and kinetic analysis of cooperative cofilin–actin filament interactions. Proc. Natl. Acad. Sci. 111, 9810–9815 (2014).

49. Hansen, S. D. et al. αE-catenin actin-binding domain alters actin filament conformation and regulates binding of nucleation and disassembly factors. Mol. Biol. Cell 24, 3710–3720 (2013).

50. Pospich, S., Merino, F. & Raunser, S. Structural effects and functional implications of phalloidin and jasplakinolide binding to actin filaments. bioRxiv 794495 (2019).

51. De La Cruz, E. M. How cofilin severs an actin filament. Biophys. Rev. 1, 51–59 (2009).

52. Sharma, S., Grintsevich, E. E., Phillips, M. L., Reisler, E. & Gimzewski, J. K. Atomic force microscopy reveals drebrin induced remodeling of F-actin with subnanometer resolution. Nano Lett. 11, 825–827 (2010).

53. Huehn, A. et al. The actin filament twist changes abruptly at boundaries between bare and cofilin-decorated segments. J. Biol. Chem. 293, 5377–5383 (2018).

54. Huehn, A. R. et al. Structures of cofilin-induced structural changes reveal local and asymmetric perturbations of actin filaments. Proc. Natl. Acad. Sci. (2020).

55. Grintsevich, E. E. et al. Mapping of drebrin binding site on F-actin. J. Mol. Biol. 398, 542–554 (2010).

56. Wilson, K. et al. Mechanisms of leading edge protrusion in interstitial migration. Nat. Commun. 4, (2013).

57. Bergert, M. et al. Force transmission during adhesion-independent migration. Nat. Cell Biol. 17, 524 (2015).

58. Martin, E. W. et al. Valence and patterning of aromatic residues determine the phase behavior of prion-like domains. Science 367, 694–699 (2020).

59. Young, K. G. & Kothary, R. Dystonin/Bpag1—a link to what? Cell Motil. Cytoskeleton 64, 897–905 (2007).

60. Davidson, P. M. et al. Actin accumulates nesprin-2 at the front of the nucleus during confined cell migration. bioRxiv 713982 (2019).

61. Fritz-Laylin, L. K. et al. Actin-based protrusions of migrating neutrophils are intrinsically lamellar and facilitate direction changes. Elife 6, e26990 (2017).

62. Nakai, N. et al. Genetically encoded orientation probes for F-actin for fluorescence polarization microscopy. Microscopy 68, 359–368 (2019).

63. Belardi, B., Hamkins-Indik, T., Harris, A. R. & Fletcher, D. A. A weak link with actin organizes tight junctions to control epithelial permeability. bioRxiv 805689 (2019).

64. Schiffhauer, E. S. et al. Mechanoaccumulative Elements of the Mammalian Actin Cytoskeleton. Curr. Biol. (2016).

65. Uyeda, T. Q., Iwadate, Y., Umeki, N., Nagasaki, A. & Yumura, S. Stretching actin filaments within cells enhances their affinity for the myosin II motor domain. PloS One 6, e26200 (2011).

66. Suzuki, N., Miyata, H., Ishiwata, S. & Kinosita Jr, K. Preparation of bead-tailed actin filaments: estimation of the torque produced by the sliding force in an in vitro motility assay. Biophys. J. 70, 401–408 (1996).

67. Sase, I., Miyata, H., Ishiwata, S. & Kinosita, K. Axial rotation of sliding actin filaments revealed by single-fluorophore imaging. Proc. Natl. Acad. Sci. 94, 5646–5650 (1997).

68. Enrique, M., Roland, J., McCullough, B. R., Blanchoin, L. & Martiel, J.-L. Origin of twist-bend coupling in actin filaments. Biophys. J. 99, 1852–1860 (2010).

69. Orlova, A. & Egelman, E. H. A conformational change in the actin subunit can change the flexibility of the actin filament. (Elsevier, 1993).

70. Phair, R. D., Gorski, S. A. & Misteli, T. Measurement of dynamic protein binding to chromatin in vivo, using photobleaching microscopy. in Methods in enzymology vol. 375 393–414 (Elsevier, 2003).

71. Stein, S. C. & Thiart, J. TrackNTrace: A simple and extendable open-source framework for developing single-molecule localization and tracking algorithms. Sci. Rep. 6, 37947 (2016).

72. Spudich, J. A. & Watt, S. The regulation of rabbit skeletal muscle contraction I. Biochemical studies of the interaction of the tropomyosin-troponin complex with actin and the proteolytic fragments of myosin. J. Biol. Chem. 246, 4866–4871 (1971).

73. Bieling, P., Telley, I. A., Hentrich, C., Piehler, J. & Surrey, T. Fluorescence microscopy assays on chemically functionalized surfaces for quantitative imaging of microtubule, motor, and+ TIP dynamics. Methods Cell Biol. 95, 555–580 (2010).

